# Parallel planning through an optimal neural subspace in motor cortex

**DOI:** 10.1101/2022.07.09.499417

**Authors:** Nicolas Meirhaeghe, Alexa Riehle, Thomas Brochier

## Abstract

How do patterns of neural activity in motor cortex contribute to the planning of a movement? A recent theory developed for single movements proposes that motor cortex acts as a dynamical system whose initial state is optimized during the preparatory phase of the movement. This theory makes important yet untested predictions about preparatory dynamics in more complex behavioral settings. Here, we analyzed preparatory activity in non-human primates planning not one, but two movements simultaneously. As predicted by the theory, we found that parallel planning was achieved by adjusting preparatory activity within an optimal subspace to an intermediate state reflecting a tradeoff between the two movements. The theory quantitatively accounted for the relationship between this intermediate state and fluctuations in the animals’ behavior down at the trial level. These results uncover a simple mechanism for planning multiple movements in parallel, and further point to motor planning as a controlled dynamical process.

## Introduction

To act efficiently, we often plan our movements ahead. Motor planning has historically been studied in experiments where subjects are given a preparatory period before executing a movement specified in advance (Day et al., 1989; Rosenbaum, 1980; Wise,1985). Behaviorally, a well-established result is that motor planning enables movements to be initiated more quickly (Crammond & Kalaska, 2000; Ghez et al., 1997; Riehle & Requin, 1989). At the neural level, the benefit of motor planning on reaction time has been linked to preparatory activity in premotor and primary motor cortex (Afshar et al.,2011; Bastian et al., 2003; Churchland & Shenoy, 2007a; Michaels et al., 2015; Riehle & Requin, 1989, 1993; Tanji & Evarts, 1976; Weinrich et al., 1984). In particular, early work has shown that preparatory activity in these brain regions is strongly modulated by key parameters (e.g., direction, speed, extent) of the upcoming movement (Churchland, Santhanam, et al., 2006; Even-Chen et al., 2019; Godschalk et al., 1985; Kurata, 1993; Messier & Kalaska, 2000; Riehle et al., 1994). These findings have led to the representational view of movement preparation which considers motor plans as parametric representations of movement features in motor cortex (Requin et al., 1991).

While the representational view provides a useful framework to describe tuning properties of motor cortical activity, it offers only limited understanding of the underlying principles governing the *dynamics* of preparatory activity (Erlhagen & Schöner, 2002). In response to this criticism, recent efforts have been made to formulate a more comprehensive theory of motor planning (Shenoy et al., 2011). The key insight was to move away from the idea of static representations at the level of individual neurons, and instead focus on the global dynamics implemented by entire neural populations in the motor cortex (Churchland et al., 2012; Fetz, 1992; Saxena & Cunningham, 2019). Following this paradigm shift, it was proposed that complex patterns of motor cortical activity ought to be viewed as a dynamical system responsible for generating the movement (Churchland et al., 2010; Sussillo et al., 2015). In this view, motor planning is seen as the process of initializing the dynamical system to the most appropriate “initial condition” to efficiently generate the desired dynamics for the movement (Churchland et al., 2010; Churchland, Yu, et al., 2006; Hennequin et al., 2014; Kao et al., 2021).

Multiple lines of evidence support this dynamical system view of motor planning. First, preparatory activity was shown to be more dynamic than previously thought (Bastian et al., 2003; Churchland, Yu, et al., 2006; Hatsopoulos et al., 2007; Rickert et al., 2009). Second, preparatory and movement-related activity can significantly differ in terms of tuning properties (Churchland et al., 2010), arguing against a purely parametric view. Third, trial-to-trial fluctuations in the preparatory state are strongly predictive of reaction times (Afshar et al., 2011; Michaels et al., 2015; Pandarinath et al., 2018; Riehle & Requin, 1993). Finally, causally perturbing the preparatory state shortly before movement initiation selectively delays movement (Churchland & Shenoy, 2007a), suggesting that preparatory activity must be in a particular state for the associated movement to be initiated (Lara et al., 2018; Vyas, O’Shea, et al., 2020).

One appeal of the dynamical systems theory is that it provides a conceptually straightforward interpretation of preparatory and motor-related signals that are notoriously difficult to parse out at the level of individual neurons (Batista et al., 2007; Churchland et al., 2012; Churchland & Shenoy, 2007b; Fetz, 1992; Scott, 2008). Indeed, the notion of initial condition has contributed to elucidating a number of computational questions at the neural population level, from the relationship between movement preparation and execution (Elsayed et al., 2016; Kaufman et al., 2014, 2015), to the role of preparatory activity in motor learning (Golub et al., 2018; Sadtler et al., 2014; Sun et al., 2020; Vyas et al., 2018; Vyas, O’Shea, et al., 2020), and the logic behind the neural control of timed movements (Remington et al., 2018; Sohn et al.,2019; Wang et al., 2017). Despite its conceptual impact on the field, however, the dynamical systems theory has remained difficult to test in experimental conditions that go beyond those that motivated its formulation. To make progress, it is thus crucial to identify and test novel predictions of the theory that can generalize to new settings.

Here we propose to focus on the case of multi-movement planning in which not one, but multiple movements are *simultaneously* prepared. Multi-movement planning poses an interesting challenge for the motor system, and can be used to expose new predictions of the initial condition (IC) hypothesis. If every movement is associated with its own IC, preparing for several potential movements should be achieved by reaching an intermediate IC located in-between the individual ICs. Further, there should exist a systematic relationship between the exact location of this intermediate state relative to individual ICs and the propensity to prepare better for one movement versus another. While prior studies have examined the neural basis of multi-movement planning (Bastian et al., 2003; Cisek & Kalaska, 2005; Dekleva et al., 2016; Ifft et al., 2012; Thura & Cisek, 2014), they were mostly concerned with assessing the coexistence of multiple motor plans in motor cortex. These studies moreover typically involved movements that were associated with different spatial locations (i.e., hand reaches toward multiple targets) and therefore could not disambiguate motor plans from the visual representations of the movement goal (Shen & Alexander, 1997; Wong & Haith, 2017). In the present study, we developed a multi-movement planning task in which monkeys had to execute one of two possible grasping movements based on a non-spatial cue. We examined the structure of preparatory activity in relation to the predictions of the dynamical systems theory, and found that our data were consistent with the initial condition hypothesis extended to the case of multi-movement planning.

## Results

### Task and behavior

Two monkeys (L and N) performed an instructed delayed reach-to-grasp task (Brochier et al., 2018; Milekovic et al., 2015; Riehle et al., 2013). The animals were trained to grasp an object using 2 possible hand grips (side grip, SG, or precision grip, PG) and subsequently pull and hold the object using 2 possible force levels (low force, LF, or high force, HF). On each trial, the animal had to wait for two successive instructions separated by a 1-s delay before initiating their movement (**Figure 1A**; Methods). The grip and force instructions were displayed via a square of 4 light-emitting diodes (LEDs) as follows: the two leftmost (resp., rightmost) LEDs instructed SG (resp., PG), while the top (resp., bottom) LEDs instructed HF (resp., LF). There were two main conditions in this task. In the “grip-cued” condition, the grip instruction was provided first, followed by the force instruction. In the “grip-uncued” condition, the force instruction was provided first, followed by the grip instruction (**Figure 1B**). In both conditions, the second instruction also served as the imperative Go signal for the animal to initiate its movement; we therefore refer to the first instruction as the “Cue”, the second instruction as the “Go”, and the 1-s delay between them as the “preparatory period”. Note that the terminology used here to describe the two task conditions differs from previous studies (Brochier et al., 2018; Milekovic et al., 2015; Riehle et al., 2013) but is more relevant to the objectives of the present study.

**Figure 1.**
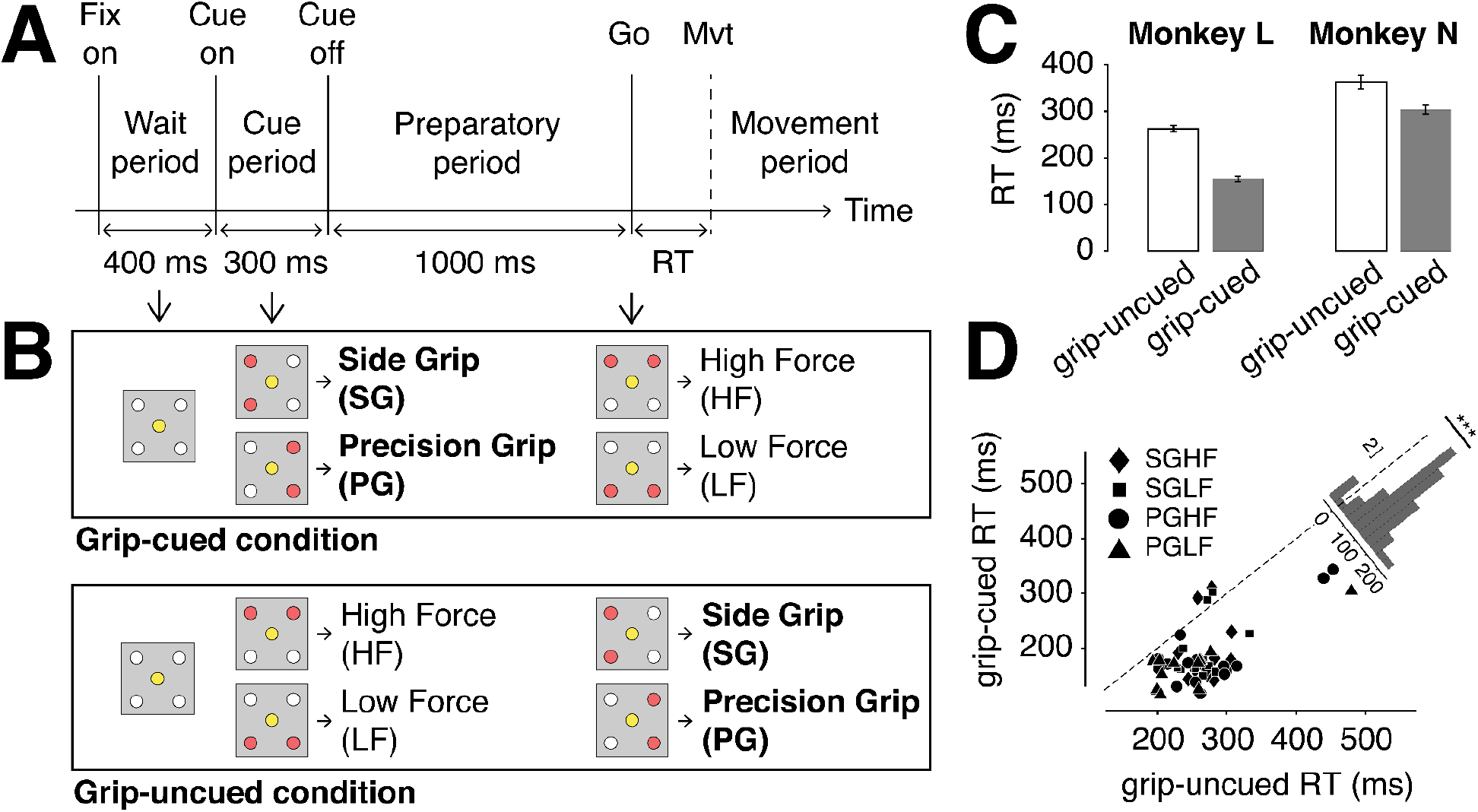
Task and behavior. **(A)** Trial structure. Each trial started with a 400-ms waiting period during which only the center LED was on. During the subsequent cue period, 2 of the 4 peripheral LEDs were illuminated for 300 ms to provide the first instruction. The cue was then turned off for 1000 ms during the preparatory period. At the end of the preparatory period, 2 other peripheral LEDs were illuminated to provide the second instruction and simultaneously signal the Go. By definition, the reaction time (RT) was defined as the time between movement initiation (vertical dashed line) and the Go signal (Methods). **(B)** Experimental conditions. In the grip-cued condition (top), the grip instruction (side grip, SG or precision grip, PG) was provided first, followed by the force instruction (high force, HF or low force, LF). In the grip-uncued condition (bottom), the force instruction was provided first, followed by the grip instruction. Instructions were provided as follows: the two leftmost (resp., rightmost) LEDs were used to instruct SG (resp., PG), and the two top (resp., bottom) LEDs were used to signal HF (resp., LF). **(C)** Reaction times for the two conditions in a typical session. In both monkeys, reaction times were shorter in the grip-cued (filled bars) compared to the grip-uncued condition (empty bars). Error bars represent 95% confidence intervals. **(D)** Average RT in the grip-cued condition plotted as a function of the average grip-uncued RT. Each point represents data from the same behavioral session. Different symbols (diamond, square, circle, triangle) indicate the trial type (resp., SGHF, SGLF, PGHF, PGLF). Inset: distribution of the difference between grip-uncued RT and grip-cued RT across sessions. This difference was significantly greater than zero (paired *t*-test, *p*<10^-10^).

In this task, when grip instruction is provided first (grip-cued), animals can plan the desired grip in advance during the preparatory period. In contrast, when grip instruction is provided last (grip-uncued), animals do not know the desired grip until the time of Go, and are therefore left uncertain about which of the two grips to plan. We used this task to study motor planning related to a single grip versus two simultaneous grips. We focused on the grip, as opposed to the force, because it is the most relevant parameter to plan the initial phase of the movement, i.e., reaching toward and grasping the object.

In sum, the task was composed of 2 conditions, with 4 trial types each (2 grips x 2 forces). The grip-cued condition was composed of PG-HF and PG-LF (hereafter collectively referred to as “PG-cued”), and SG-HF and SG-LF (“SG-cued”) trials, while the grip-uncued condition was composed of HF-PG and LF-PG (“PG-uncued”) and HF-SG and LF-SG (“SG-uncued”) trials. Note that the 4 trial types were identical across conditions in terms of final movement, but differed only in the order that the grip/force information was provided. The 4 trial types were randomly interleaved within each condition, and the conditions were performed in separate blocks of trials within the same behavioral session (Methods).

At the end of training, animals were proficient in all conditions and trial types (average success rate: 92% for monkey L; 97% for monkey N; **Table S1**). These high success rates indicate that the monkeys were able to interpret and use the instruction cues to rapidly alternate between the desired actions. Critically, we confirmed that animals used the grip information (when available) to plan their movement ahead. Indeed, reaction times (RT; defined between the time of Go and movement initiation) were shorter in the grip-cued compared to the grip-uncued condition (RT = mean+sem; RT_cued_ = 155±3 ms, RT_uncued_ = 263±4 ms, *p*<10^-10^ for monkey L; RT_cued_ = 303±5 ms, RT_uncued_ = 363±8 ms, *p*<10^-9^ for monkey N; ANOVA on RT testing for the main effect of grip-cued versus grip-uncued; **Figure 1C**), and this effect was robustly observed across animals and behavioral sessions (paired *t*-test on across-session RT_cued_ versus RT_uncued_, *p*<10^-10^; **Figure 1D**). These results are in line with numerous previous studies showing a beneficial effect of movement preparation on reaction time (Ames et al., 2014; Churchland & Shenoy, 2007a; Crammond & Kalaska, 2000; Ghez et al., 1997; Riehle,2005; Riehle & Requin, 1989, 1993; Zaepffel & Brochier, 2012).

### Single-neuron activity underlying single and dual grip planning

To investigate the neural basis of motor planning in this task, we analyzed spiking activity in the primary (M1) and premotor (PM) cortex recorded via chronically-implanted multi-channel arrays (Brochier et al., 2018). Prior studies using this task have explored how the spatio-temporal structure of local field potentials and spiking activity in PM/M1 relate to various aspects of animals’ behavior (Denker et al., 2018; Milekovic et al.,2015; Riehle et al., 2013, 2018; Torre et al., 2016). None of these studies have raised the question of multi-grip planning. Here, we sought to directly compare preparatory activity in the grip-uncued and grip-cued conditions to relate the planning process of two potential grips to the planning of a single grip.

Our epoch of interest included the 300-ms presentation of the Cue (providing grip information in grip-cued, and force information in grip-uncued), followed by the 1-s preparatory period preceding the Go signal, and the subsequent movement period. In this epoch, single neurons had widely varied activity profiles (**Figure 2A, red and blue lines**). In the grip-cued condition, some neurons encoded the grip type shortly after the Cue (**Figure 2A (iii)**), or right before the Go (**Figure 2A (ii)**), while others remained sensitive to the grip throughout the preparatory period (**Figure 2A (i)**). At the population level, grip tuning depth (measured as the instantaneous firing rate difference between PG-cued and SG-cued) was heavily distributed across neurons and highly dynamic across time points and epochs (**Figure 2B, S1**). Moreover, tuning properties during movement preparation and execution were largely unrelated (**Figure 2C**), in line with previous reports (Churchland et al., 2010; Elsayed et al., 2016).

**Figure 2.**
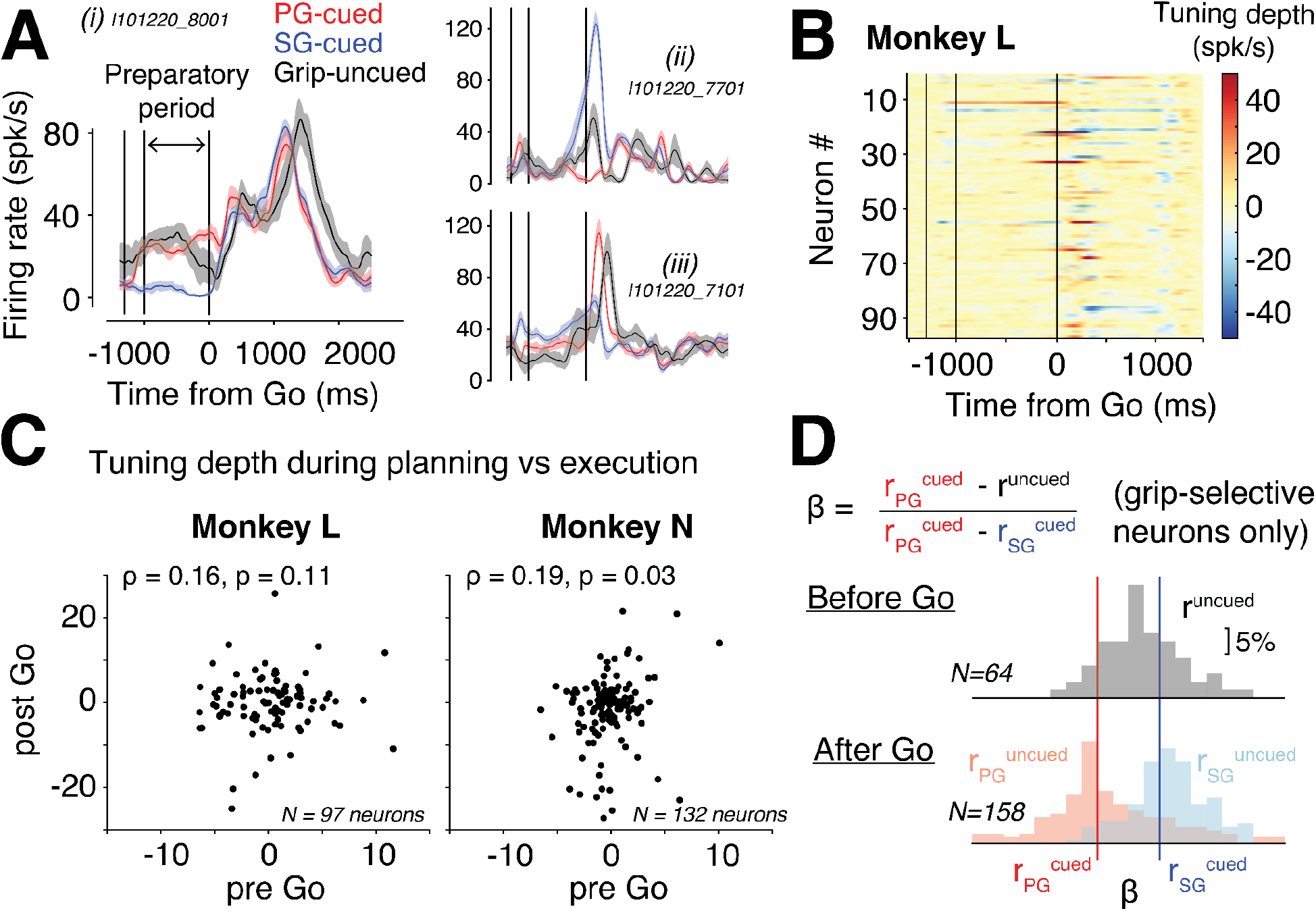
Neural encoding of grip information. **(A)** Firing rate of four example neurons from the two monkeys aligned to the Go signal. Neural activity is color-coded by condition; red for PG-cued (here, PGHF), blue for SG-cued (SGHF), and black for grip-uncued (HFPG). Firing rates were obtained by binning (bin = 20ms) and smoothing (SD of Gassian kernel = 40 ms) spike counts averaged across trials of the same condition. Shaded areas denote 99% CI obtained from bootstrap resampling of trials (N=100). Vertical lines represent Cue on, Cue off, and Go. **(B)** Grip tuning depth across neurons, as measured by the instantaneous firing rate difference between the PG-cued and SG-cued activity (we used the same time resolution as in (A) for the firing rates). Each row represents one neuron, and the color indicates the magnitude of the tuning depth (blue for negative values, i.e., neuron has a larger firing rate in SG; red for positive values). See Figure S1A for the second monkey. **(C)** Correlation between tuning depths measured in a 20-ms window 500 ms before and after the Go signal. In both monkeys (left and right), the Pearson correlation was not significantly different from zero (p>0.01). The number indicated in italic is the number of neurons considered. **(D)** Comparison of firing rates between the grip-cued and grip-uncued condition. For each grip-selective neuron (i.e., neuron whose preparatory activity significantly differed between PG-cued and SG-cued; see Methods), we computed the ratio 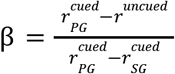, where 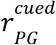, 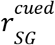 and *r^uncued^* designate the firing rate of the neuron respectively in the PG-cued, SG-cued, and grip-uncued condition. By definition, β is close to 0 (red vertical line) if *r^uncued^* is close to 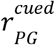, and close to 1 if *r^uncued^* is close to 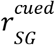. Top: the black histogram represents the distribution of β across grip-selective neurons when computing firing rates in a 20-ms window 200 ms before Go. Bottom: distributions of β across grip-selective neurons when computing firing rates in a 20-ms window 400 ms after Go. The distributions are color-coded based on the grip revealed at Go (pale red for PG-uncued; pale blue for SG-uncued). To increase the number of grip-selective neurons considered (indicated in italic), data from both monkeys and force levels were combined.

In the grip-uncued condition, single neurons had complex and highly heterogeneous activity profiles (**Figure 2A, black lines**). During the preparatory period, many neurons reached a level of activity before Go that was different from that associated with the PG-cued or SG-cued condition (**Figure 2D**). While some neurons appeared to reach an intermediate level of activity between their PG-cued and SG-cued activity levels (e.g., neuron (ii) in **Figure 2A**), we found that this was not systematic at the population level: a sizable fraction of grip-selective neurons (45% for monkey L; 39% for monkey N; averaged across grip-uncued condition; Methods) had firing rates outside of their grip-cued range.

Together, these observations are in keeping with prior studies showing that motor cortex is tuned to upcoming movement parameters (Churchland, Santhanam, et al., 2006; Even-Chen et al., 2019; Godschalk et al., 1985; Kurata, 1993; Messier & Kalaska, 2000; Riehle et al., 1994), i.e., here the grip type. However, they also demonstrate a complex relationship at the single-neuron level between grip-cued and grip-uncued activity, which are difficult to explain in the context of classic tuning-based representational analyses.

### The optimal subspace hypothesis for multi-movement planning

To parse out the relationship between grip-cued and grip-uncued preparatory activity, we leveraged a recent theory of motor planning rooted in the framework of dynamical systems (Churchland et al., 2010; Hennequin et al., 2014; Kao et al., 2021; Vyas, Golub, et al., 2020). According to the “initial condition” (IC) hypothesis, preparatory activity should be viewed as a dynamic optimization process evolving toward an optimal state from which movement-related dynamics can be efficiently generated (Afshar et al.,2011; Ames et al., 2014; Hennequin et al., 2014; Kao et al., 2021). Consistent with this view, we found that population dynamics in the two grip-cued conditions rapidly diverged following the grip cue presentation, and remained separated throughout the preparatory period (**Figure S2A**), which may reflect the evolution toward two grip-specific initial conditions. In comparison, the two force-specific trajectories in the grip-uncued condition did not diverge as significantly during the preparatory period (**Figure S2A**).

Although the IC hypothesis was originally introduced for single movements, one may extend it to the case of multi-movement planning. If each movement is associated with its own initial condition (single-movement IC), we can formulate two predictions: the preparatory state associated with multiple alternatives should 1) lie within the subspace containing all the single-movement ICs, and 2) be located “in-between” the single-movement ICs to rapidly converge to one of them when the desired movement is finally prescribed. We refer to this augmented view of the IC hypothesis as the “optimal subspace hypothesis”. Note that this term has previously been introduced in the context of single planned movements (Churchland, Yu, et al., 2006), but has a different meaning here, since it is used to extend the IC hypothesis to multi-movement planning.

To test the predictions of the optimal subspace hypothesis, we turned to population-level analyses, by considering population activity as a collection of states (i.e., neural trajectory) evolving in a high-dimensional space where each dimension represents the activity of one neuron (Sohn et al., 2020). In this state space, we defined the subspace containing the preparatory states associated with each grip. Since there were only two grips, this subspace was 1-D, and defined by the vector connecting the SG-cued IC to the PG-cued IC (**Figure 3A**); we refer to this dimension as the “grip dimension”. Note that because the two grip-cued trajectories did not remain parallel throughout the preparatory period (**Figure S2A**), the grip dimension was defined at every time point (Methods). We then projected both the grip-cued and grip-uncued trajectories onto the grip dimension to evaluate the proximity of the grip-uncued state relative to the two grip-cued states as a function of time in the trial (**Figure 3A and S3C**).

**Figure 3.**
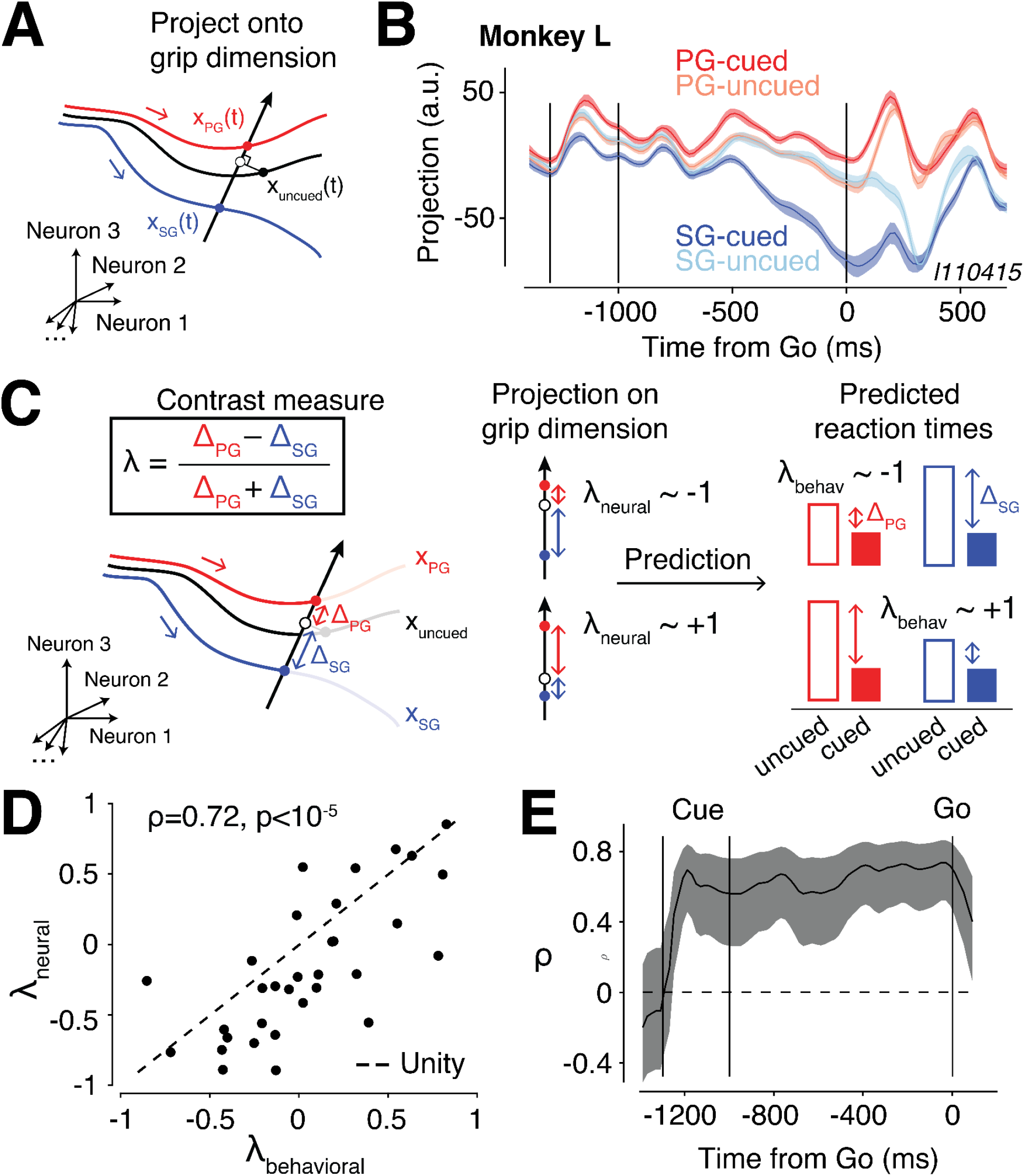
Test of the optimal subspace hypothesis. **(A)** Schematic of the projection onto the grip dimension (black vector) in the state space. At any time point, the grip dimension was defined as the unit vector pointing from the SG-cued state (*x_SG_*(*t*), filled blue circle) to the (*x_PG_*(*t*) filled red circle). The schematic shows the orthogonal projection (open black circle) of the grip-uncued state (*x_uncued_*(*t*), filled black circle) onto the grip dimension. **(B)** Temporal evolution of the projections onto the grip dimension within the preparatory and movement epochs. The dark red and blue represent the projections of the PG-cued and SG-cued trajectories, respectively. The projection of the grip-uncued trajectories is shown separately for the PG-uncued (pale red) and the SG-uncued (pale blue) condition. Data from one session is shown; see Figure S2B for more sessions. The slight separation of PG-cued and SG-cued projections before the Cue onset is an artifact of defining the grip dimension in a non-cross-validated way (which is not necessary for this particular analysis, since we are interested in projecting grip-uncued trials, which by definition are cross-validated). Note however that a cross-validated neural distance analysis confirmed that the PG-cued and SG-cued trajectories overlapped prior to Cue onset (Figure S2A). **(C)** Predictions of the optimal subspace hypothesis. Left: to evaluate the relative positions of the PG-cued, SG-cued and grip-uncued preparatory states along the grip dimension, we defined for each experimental session a contrast measure (λ_*neural*_) as follows: 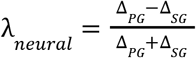, where Δ_*PG*_ and Δ_*SG*_ represented the distance of the grip-uncued projection to the PG-cued and SG-cued state, respectively. Middle: λ_*neural*_ close to −1 (top) corresponded to the grip-uncued projection close to the PG-cued state; λ_*neural*_ close to +1 (bottom) corresponded to the grip-uncued projection close to the SG-cued state. Right: to assess the influence of the preparatory state on behavior, we defined another contrast measure (λ_*behavioral*_) using the same expression as above, except Δ_*PG*_ and Δ_*SG*_ now represented the difference in reaction time between the grip-uncued and grip-cued condition for PG and SG, respectively. The optimal subspace hypothesis predicted that λ_*neural*_ and λ_*behavioral*_ should correlate. **(D)** Correlation between λ_*neural*_ and λ_*behavioral*_. Each dot represents one session. Sessions were pooled across animals. The dashed line shows the unity line. **(E)** Pearson correlation between λ_*neural*_ and λ_*behavioral*_ as a function of time in the preparatory period. The shaded area denotes the 95% confidence interval (Methods).

Before the Cue presentation, the preparatory states largely overlapped (**Figure 3B**; but see **Figure S2A** for a cross-validated neural distance analysis of the trajectories), which was expected since no grip information was available at that time. Following the Cue, the two grip-cued states rapidly diverged and remained separated throughout the preparatory and movement period, consistent with our previous distance analysis (**Figure S2A**). The grip-uncued state had a qualitatively similar temporal profile, but notably remained *in-between* the two grip-cued states until ~100 ms after Go (**Figure 3B**). The trajectory then separated from this intermediate state into two grip-uncued trajectories (PG-uncued and SG-uncued) which rapidly merged with the corresponding grip-cued trajectories (PG-cued and SG-cued, respectively) about 200-250 ms after Go. The same observations were made in the two animals, and confirmed using both force levels of the grip-uncued condition (**Figure S2B**).

We verified that the intermediate state in the grip-uncued condition did not simply reflect an absence of change from baseline activity: the euclidean distance between neural states immediately before and after the Cue was statistically different from zero (95% CI = [0.03 0.04] for HF, and [0.03 0.04] for LF in monkey L; [0.02 0.03] for HF, and [0.02 0.03] for LF in monkey N), indicating a significant change from baseline similar to that found in the grip-cued condition (95% CI = [0.05 0.06] for PG, [0.05 0.06] for SG in monkey L; [0.03 0.04] for PG, [0.03 0.04] for SG in monkey N). This analysis demonstrates that, before Go, the grip-uncued activity evolved toward an intermediate preparatory state along the dimension that contained the two grip-cued preparatory states, and rapidly converged to one of them once grip information was provided.

### Optimization of initial conditions explains inter-session variability

So far, our results are consistent with the optimal subspace hypothesis: preparatory activity in the grip-uncued condition reaches an intermediate state between the two grip-specific initial conditions. This intermediate state might result from an optimization process facilitating the execution of the two potential grips. To firmly establish this result, however, we ought to demonstrate a tighter relationship between the preparatory state and the animal’s behavior. In particular, the exact position of the grip-uncued initial condition along the grip dimension should be predictive of animals’ tendency to favor the preparation of one grip versus the other. For instance, if the grip-uncued IC is slightly closer to one of the two grip-cued ICs, say PG-cued (as in **Figure 3B**), then this should confer a slight reaction time benefit to PG relative to SG. In the following, we therefore sought to leverage the inter-session variability in the neural and behavioral data to test this stronger prediction of the optimal subspace hypothesis.

To quantify the proximity of the grip-uncued IC relative to the two grip-cued ICs along the grip dimension for each session, we defined a neural contrast, 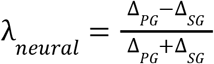, where Δ_*PG*_ and Δ_*SG*_ respectively represent the distance of the grip-uncued projection to the PG-cued and SG-cued state along the grip dimension. By definition, λ, was λ_*neural*_ bounded by −1 and 1, and was negative if the grip-uncued IC was closer to the PG-cued IC, and positive if the grip-uncued IC was closer to the SG-cued IC (**Figure 3C**). According to the optimal subspace hypothesis, λ_*neural*_ closer to −1 should confer a slight advantage to the planning of PG relative to SG (**Figure 3C, middle top**). That is, there should be a smaller RT difference between PG-uncued and PG-cued trials compared to SG-uncued and SG-cued trials (**Figure 3C, top right**). Conversely, λ_*neural*_ closer to +1 should confer a slight advantage to SG over PG (**Figure 3C, middle bottom**), with a smaller RT difference between SG-uncued and SG-cued compared to PG-uncued and PG-cued (**Figure 3C, bottom right**). To assess this effect in behavior, we defined a behavioral contrast, 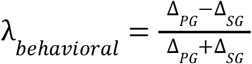, where Δ_*PG*_ and Δ_*SG*_ respectively represent the (absolute) reaction time difference between the grip-uncued and grip-cued condition for PG and SG. By definition, λ_*behavioral*_, was close to −1 if the RT difference between the cued and uncued condition was smaller for PG than SG, and +1 if the RT difference was smaller for SG than PG.

Based on the optimal subspace hypothesis, we expected λ_*neural*_ and λ_*behavioral*_ to covary. Consistent with this prediction, we found a strong positive correlation between the two contrast measures across sessions (Pearson correlation, *ρ*=0.72; *p*<10^-5^; **Figure 3D**). This effect was specific to the grip dimension, since the correlation was abolished when we projected the preparatory states onto a dimension orthogonal to the grip dimension but constrained to capture a sizable amount of the variance in the data (*ρ*=0.13; *p*=0.48; see Methods). When we extended our correlation analysis using neural activity at various time points of the cue and preparatory periods, we found that the correlation between λ_*neural*_ and λ_*behavioral*_ became positive around 100ms after the Cue presentation (**Figure 3E**). Moreover, the correlation was strongest when λ_*neural*_ was computed right at the time of Go, i.e., at the time most relevant for setting up the initial condition for the upcoming movement. Together, these results provide compelling evidence that the preparatory state in the grip-uncued condition was optimized to fall in-between the two grip-cued initial conditions, and that the extent to which this optimization favored one grip over the other was quantitatively reflected in the animals’ behavior across sessions.

### The optimal subspace hypothesis holds at the single-trial level

Our previous analysis was based on trial-averaged activity, and did not take into account the variability that occurred across trials of a given session. As a final test of the optimal subspace hypothesis, we sought to verify its predictions down at the single-trial level. One important assumption of the optimal subspace hypothesis is that neural trajectories reach a fixed state (i.e., “threshold”) when the movement is triggered, and that the distance between the initial state (at the time of Go) and the threshold is what determines the reaction time. As a first step, we thus examined neural dynamics leading up to movement initiation to test for the presence of a putative threshold common across conditions. We aligned neural activity to movement onset, and found that dynamics indeed reached a systematic level of activity at movement initiation, independent of the condition type (cued versus uncued; **Figure S5**). Building on this result, we next sought to explain the trial-by-trial differences in reaction times based on the initial distance of the preparatory state to the threshold.

Let us consider a particular trial in which the projection of the grip-uncued preparatory state along the grip dimension is slightly biased toward the PG-cued IC (**Figure 4A**). Since we (arbitrarily) defined the grip dimension as pointing from the SG-cued IC to the PG-cued IC, this bias corresponds to a *large* projection onto the grip dimension. According to the optimal subspace hypothesis, this bias should provide a “head-start” to the execution of PG, and thus have a *beneficial* effect on the reaction time when the grip type revealed at the time of Go is PG, i.e., in the PG-uncued condition. As a result, the optimal subspace hypothesis predicts a *negative* correlation between trial-by-trial projections and RTs in the PG-uncued condition, i.e., *larger* projections onto the grip dimension lead to *shorter* reaction times (**Figure 4B, top**). The hypothesis predicts the opposite effect in the SG-uncued condition: there should be a *positive* correlation between trial-by-trial projections and RTs, i.e., *larger* projections onto the grip dimension lead to *longer* reaction times (**Figure 4B, bottom**).

**Figure 4.**
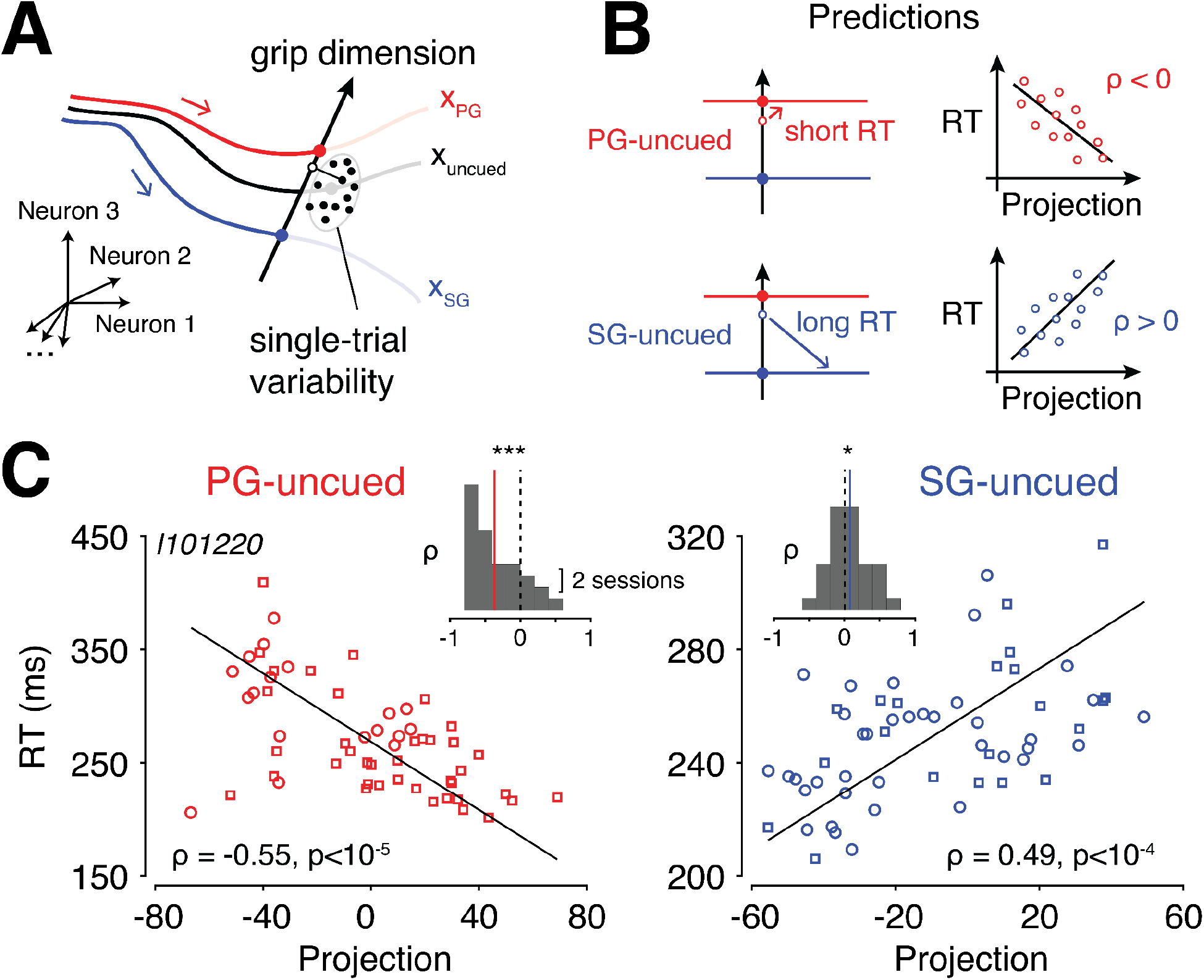
Predicting trial-by-trial variability. **(A)** Schematic of single-trial projections onto the grip dimension. Large filled circles represent trial-averaged states, while smaller circles represent individual trials. **(B)** Single-trial predictions of the optimal subspace hypothesis. One particular grip-uncued trial biased toward the PG-cued state is shown. If the grip revealed at Go is PG (i.e., PG-uncued condition, in red at the top), the reaction time (RT) for this trial is expected to be short. This predicts a negative correlation between RT and the projection (right). In contrast, if the grip revealed at Go is SG (i.e., SG-uncued condition, in blue at the bottom), the reaction time for this trial is expected to be long. This predicts a positive correlation between RT and the projection. **(C)** Correlation between RT and single-trial projections for the PG-uncued (left) and SG-uncued (right) condition in one high-yield session. Individual trials corresponding to the two force levels are shown with different symbols (circle for HF; square for LF) and combined to increase statistical power. Inset: Summary of correlation values across sessions (*p<0.05, ***p<0.001).

To test these predictions, we estimated single-trial neural dynamics using a 200-ms sliding window, and projected individual trajectories onto the grip dimension (Methods). Because estimating single-trial activity is prone to noise, we focused on a high-yield session with a large number (n=97) of simultaneously recorded neurons, and a large number (n=141) of successful trials (see **Table S1**). Trial-by-trial preparatory activity appeared to be highly variable along the grip dimension, particularly near the time of Go, and the amount of trial-by-trial variance accounted for by the grip dimension was approximately 10 times larger than expected by chance (**Figure S3A**). For each individual trial, we plotted the neural projection immediately prior to Go as a function of the reaction time, separately for the PG-uncued and SG-uncued conditions (**Figure 4C**). As predicted by the optimal subspace hypothesis, we found a negative correlation in the

PG-uncued condition (Pearson correlation, *ρ_PG_*=-0.55; *p*<10^-5^), and a positive correlation in the SG-uncued condition (*ρ_SG_*=0.49; *p*<10^-4^). This result held across recorded sessions, although the effect was weaker likely due to the lower number of neurons in these sessions (one-sided *t*-test on *ρ* across sessions, mean+sem, *ρ_PG_*=-0.37±0.06; *t*(31)=-5.97, *p*<10^-6^; *ρ_SG_*=0.09±0.05, *t*(31)=1.73, *p*<0.05; **Figure 4C, inset**). Moreover, the same patterns of correlation between RTs and trial-by-trial projections were found for the grip-cued conditions (**Figure S3B**), reinforcing the idea that the grip dimension was key in controlling the initial condition for planning the movements. Altogether, these results confirm the tight relationship predicted by the optimal subspace hypothesis between deviations of the preparatory state along the grip dimension and animals’ reaction times down at the single-trial level.

### Ruling out rapid vacillations in preparatory dynamics

Our previous analysis shows that trial-by-trial variations in reaction times can be predicted from variations in the exact location of the preparatory state along the grip dimension at the time of Go. One remaining question is to identify the source of this variability. Previous studies have shown that when instructed movements are not fully prescribed in advance, or are likely to change during the planning phase of the movement, preparatory activity can rapidly *vacillate* between the preparatory states associated with the various possibilities (Kaufman et al., 2015; Peixoto et al., 2021). This phenomenon could in principle explain the correlation we observed between neural projections and reaction times, in that RT would depend on how complete the transition toward one of the cued states is at the time of Go (Dekleva et al., 2018).

In the following, we sought to test this possibility, which we refer to as the “vacillation hypothesis”, against the alternative hypothesis of a stable preparatory state throughout the preparatory period (“stable preparation hypothesis”). First, we assessed the extent to which uncued preparatory activity oscillated between the two cued states by looking at cross-temporal correlations. One direct prediction of the vacillation hypothesis is that the projection of the uncued state onto the grip dimension might be close to the PG-cued state at one time point of the trial, and become closer to the SG-cued state at some later time point. This predicts a *negative* correlation between single-trial projections considered at different time points far enough apart during the preparatory period: if the projection is large early on (i.e., close to the PG-cued state), it will be small later on (i.e., close to the SG-cued state), and vice-versa. Alternatively, the stable preparation hypothesis posits that the proximity of the uncued state relative to the two cued states should be maintained throughout the trial, which predicts a *positive* correlation across time.

To test these predictions, we calculated the correlation between single-trial projections computed at different time lags within the pre-cue, preparatory, and movement periods. For this cross-temporal correlation analysis, we used non-smoothed spiking data (binned at a 100-ms resolution) to avoid creating spurious correlations in time. We found that pre-cue projections became rapidly uncorrelated with projections during the preparatory period (**Figure 5A**), which was expected since animals have no information prior to the Cue. During the preparatory period, however, a clear pattern appeared where projections remain positively correlated (correlation above 0.3) regardless of the time lag between the projections (**Figure 5A**). This result shows that single-trial projections which are close to a particular cued state at one point of the preparatory period remain on average close to that state throughout the preparatory period, which argues against the vacillations hypothesis and in favor of a stable preparatory state.

**Figure 5.**
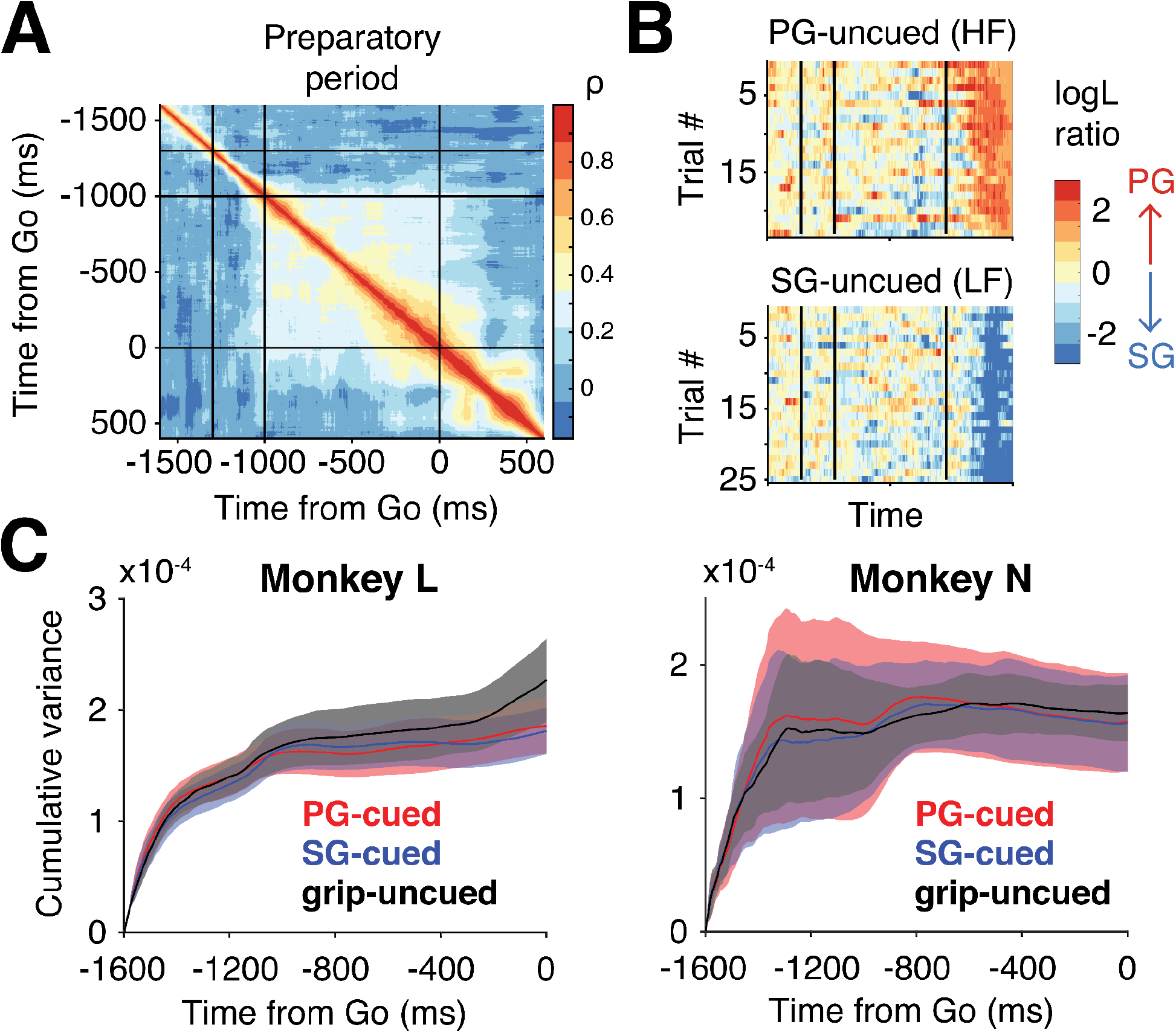
Test of the vacillation hypothesis against the stable preparation hypothesis. **(A)** Cross-correlation of grip-uncued single-trial projections along the grip dimension. The result is shown as a heatmap where each pixel represents the correlation between projections computed at two different time points of the trial (*t_1_* and *t_2_*; abscissa and ordinate of the heatmap). We gathered projections computed at *t_1_* and *t_2_* into two separate vectors of size [nTrialx1] and calculated the Pearson correlation between the two vectors. For this analysis, to increase statistical power, we normalized the grip-uncued projections within each session by the associated grip-cued projections to allow pooling the data across sessions and animals. The data shown is from the PG-uncued (HF) condition. **(B)** Log likelihood ratio of single-trial projections in the grip-uncued condition. At every time point, we modeled the distributions of single-trial projections along the grip dimension of the PG-cued and SG-cued conditions as Gaussian distributions. We then computed the likelihood that, at every time point, each grip-uncued trial comes from either the PG-cued or SG-cued distribution. To interpret the result, we further computed the ratio of these likelihoods, and took the log_10_ of this ratio. A log likelihood ratio of 0 indicates that a given trial is equally likely to come from the PG-cued or the SG-cued condition, while a log likelihood ratio of +2 indicates that a given trial is 100 times more likely to come from the PG-cued condition. **(C)** Evolution of the cumulative variance of single-trial projections along the grip dimension. The variance was computed after subtracting the mean trajectory within each condition. Colors indicate the condition, grip-cued (red for PG, blue for SG) and grip-uncued (black). Shaded areas denote the 95% confidence interval computed as 1.96 times the SEM across sessions.

One remaining concern is that a variable timescale of the vacillations across trials may obfuscate the result of the previous analysis. For instance, if some trials vacillate more slowly than others (in the extreme case, there may be no vacillation at all in some trials), then the correlation across trials could still remain positive despite the presence of vacillations. To confirm our result, we thus performed a second analysis focusing on each individual trial and directly inspired by the method developed in (Kaufman et al.,2015). Specifically, we projected single-trial neural states of the grip-cued and grip-uncued conditions onto the grip dimension at every time point of the trial. We then modeled the PG-cued and SG-cued single-trial distributions by two Gaussians, and computed the likelihood that, at any given time, the uncued preparatory state belonged to either the PG-cued or SG-cued distribution (Methods).

As can be seen in **Figure 5B**, the majority of single-trial uncued trajectories appeared to show a consistent bias for one or the other cued state during the preparatory period, while some trials do exhibit more variations in time. To determine whether these fluctuations represent purposeful vacillations between the two cued states, or instead result from private spiking variability at the level of single trials, we sought to estimate the strength of these fluctuations and compare it to some baseline. To do so, we computed the variance across time points of the neural projections along the grip dimension; if fluctuations occur during the preparatory period, this variance should be large. As a baseline, we computed this variance for the grip-cued condition in which animals know which grip to prepare, so that any fluctuations can be considered baseline variability. We then compared this variance to the variance associated with the two grip-uncued conditions. We found that, in both monkeys, the variance was similar between the grip-cued and grip-uncued conditions (**Figure 5C**). This result indicates that grip-uncued trajectories did not exhibit more excursions away from the mean trajectory than expected from baseline variability.

Together, these two analyses argue against the vacillations hypothesis, and support instead the stable preparation hypothesis according to which preparatory activity is maintained to a stable intermediate preparatory state during planning.

## Discussion

In this study, we analyzed preparatory activity in the motor cortex of monkeys planning two movements in parallel. Our results supported an augmented view of the initial condition hypothesis inspired by the theory of dynamical systems (Afshar et al., 2011; Churchland et al., 2010; Churchland, Yu, et al., 2006; Shenoy et al., 2013). We found that the preparatory state associated with two possible movements lies within an optimal subspace containing the preparatory states associated with each movement planned separately. This intermediate state serves as a suitable initial condition to rapidly reconfigure motor cortical dynamics for the execution of the two movements.

In light of prior studies suggesting different roles of primary motor (M1) and premotor (PM) cortex in movement preparation (Churchland & Shenoy, 2007a; Crammond & Kalaska, 2000), we examined this effect in both structures, and found that the tight relationship between preparatory activity and reaction times held when considering putative PM and M1 neurons separately (**Figure S4**). We note however that our classification was coarse and limited by differences in the exact recording sites across animals. We therefore cannot exclude functional differences in relation to parallel planning at a finer anatomical scale between premotor and primary motor cortex.

A number of previous studies on multi-movement planning have reported concurrent representations of motor plans (Bastian et al., 2003; Cisek & Kalaska, 2005; Coallier et al., 2015; Dekleva et al., 2016; Klaes et al., 2011; Thura & Cisek, 2014). These studies typically involved visually-guided movements that were associated with different spatial locations (i.e., hand reaches toward multiple potential targets). It is therefore possible that preparatory activity observed in these tasks reflected a spatial/directional tuning to the targets (Cisek, 2012; Shen & Alexander, 1997), which may have contributed to biasing the results toward a representational view (Cisek, 2006, 2007). Other studies employing a task similar to ours did find evidence for the co-activation of neurons tuned to different grip types (Baumann et al., 2009; Fluet et al., 2010). This result is not necessarily at odds with ours. Indeed, we did find that grip-selective neurons were active in the preparatory period of the grip-uncued condition. However, we showed that the activity level reached by grip-selective neurons did not reflect either one of the two levels associated with the grip-cued condition.

Our neural findings shed new light on a large body of behavioral studies showing an “averaging effect” of pre-planned movements when faced with two possible alternatives (Arai et al., 2004; Chapman et al., 2010; Chou et al., 1999; Gallivan & Chapman, 2014; Ghez et al., 1997; Hudson et al., 2007; Stewart et al., 2014). This effect has typically been attributed to two concurrent motor plans competing and blending during movement execution (Cisek, 2007; Stewart et al., 2013). Our results offer a different explanation: rather than representing the two motor plans in parallel, the motor cortex may select a single preparatory state which achieves a tradeoff between the two possible movements and optimizes task performance (Alhussein & Smith, 2021; Costello et al., 2013; Gallivan et al., 2015; Haith et al., 2015; Wong & Haith, 2017). In the case of reaching movements (as often used in prior studies), because target directions are parametrically organized in space, this intermediate state may naturally lead to a reach aimed at the average of the two potential targets.

It could perhaps be argued from a representational standpoint that the two grips are jointly encoded at a single-neuron level by setting firing rates in the uncued condition to an intermediate level between PG-cued and SG-cued activity. The issue with this interpretation, however, is that the representational view does not directly prescribe what this intermediate level of activity should be for each neuron. One possibility that we empirically tested and rejected is the idea that firing rates in the grip-uncued condition should all be at the average of their PG-cued and SG-cued activity levels. Our results demonstrate, by contrast, the power of the dynamical systems approach to explain behavioral and neural observations at the population-level. Crucially, predictions of the dynamical systems view apply to the population as a whole, and not to individual neurons. That is, single neurons do not necessarily need to reach a specific level of activity and follow a consistent rule across neurons. Moreover, the dynamical systems view predicts a clear relationship between preparatory states and reaction times, which is not directly predicted by the representational view. Nevertheless, we believe that both frameworks can and should perhaps co-exist, albeit at different levels of description.

Could the intermediate initial condition reflect an intermediate grip between PG and SG which only diverges mid-flight into either PG or SG? Several lines of evidence argue against this possibility. First of all, grasping movements––unlike reaching movements which are classically used in multi-movement planning tasks––are not represented on a continuum but rather correspond to discrete movement categories. As a result, it is unclear what an intermediate grip between two categorically-distinct grasping movements means. Second, the large across-session variability of the intermediate state also argues against the possibility of animals preparing a specific intermediate movement, which would result in a well defined and less variable preparatory state. Finally, if the animals were planning an intermediate movement, we would not find a difference in reaction time between the two uncued conditions, i.e., when PG (PG-uncued) or SG (SG-uncued) is revealed at the time of Go. For these reasons, we believe that the intermediate preparatory state in the grip-uncued condition is unlikely to reflect the planning of an intermediate movement.

Previous studies reported that preparatory dynamics can rapidly “vacillate” between multiple states when the instructed movement is not prescribed in advance (Kaufman et al., 2015; Peixoto et al., 2021). We have specifically tested this hypothesis, and ruled out the possibility that preparatory activity vacillates between the two preparatory states associated with the two grips. One possible reason for why we did not find vacillations is that our experimental design did not involve a dynamically changing movement instruction, e.g., virtual barriers that could move/appear at any time during movement preparation, which may incentivize animals to exhibit covert “changes of mind”. Whether vacillations of the intermediate state in the grip-uncued condition could be induced by manipulating the uncertainty about the upcoming grip remains to be investigated.

To our knowledge, this study constitutes the first validation of the initial condition hypothesis in the context of multi-movement planning. Previous studies were mostly restricted to single planned movements (Afshar et al., 2011; Elsayed et al., 2016; Even-Chen et al., 2019; Kaufman et al., 2014; Vyas et al., 2018), or multiple movements planned in rapid sequence (Ames et al., 2014, 2019; Zimnik & Churchland, 2021). By generalizing the notion of initial condition to movement preparation under uncertainty, our results reinforce the idea that motor planning should be seen as a dynamical process optimized to generate appropriate movement-related dynamics (Churchland & Shenoy, 2007a; Kao et al., 2021). Although the anatomical substrate underlying this optimization process is beyond the scope of our study, the thalamo-basal ganglia-cortical loop is a natural candidate (Athalye et al., 2020; Kao et al., 2021). Low-dimensional inputs to the motor cortex (Dubreuil et al., 2022; Logiaco et al., 2021; Sauerbrei et al., 2020) could for instance serve to adjust the initial state within the optimal subspace dimensions (Beiran et al., 2021; Sohn et al., 2020). Further investigations will be needed to elucidate this circuit-level question.

Our study investigated motor planning associated with only two movements. While we cannot claim that our results will generalize, we can formulate a testable prediction for cases with more alternatives. For N possible movements, we predict that preparatory activity will be located in the state space so as to minimize the distance to the N initial conditions. This optimization may be facilitated if movements are naturally organized along a parametric continuum (e.g., reaching directions); in this case, the appropriate IC may correspond to one existing IC located in-between the others. For non-parametric movements, it may be more challenging for motor cortex to find the appropriate initial condition, which would result in larger variability in the position of the intermediate IC (as seen in this study). Future studies could also test whether one movement being more likely than the others systematically biases the location of the preparatory state toward the associated initial condition (Dekleva et al., 2018), or more generally, if each initial condition is weighted by the probability that each movement will be executed.

## Methods

### Experimental procedures

All procedures were approved by the local ethical committee (C2EA 71; authorization A1/10/12) and conformed to the European and French government regulations. Experiments involved two naive, awake, behaving monkeys (species: *Macaca mulatta*; ID: L and N; sex: female and male; weight: 6 and 7 kg; age: 6 years old). Monkeys were implanted with a 96-channel chronic Utah array (Blackrock Microsystems, Salt Lake City, UT, USA) in the motor cortex contralateral to the working hand (right hemisphere for both monkeys). The exact array location can be found in (Brochier et al., 2018) and in Figure S4. Data were recorded using the 128-channel Cerebus acquisition system (Blackrock Microsystems, Salt Lake City, UT, USA). Analysis of behavioral and neural data was performed using MATLAB (Mathworks, MA).

### Behavioral task

Two monkeys performed an instructed delayed reach-to-grasp-and-pull task previously described in (Riehle et al., 2013). Briefly, the animals sat in a primate chair opened at the front and were trained to grasp an object (stainless steel parallelepiped, 40 mm x 16 mm x 10 mm, angled at 45° from the vertical and located about 20 cm away from the monkey) using 2 possible hand grips (side grip, SG, and precision grip, PG) and subsequently pull and hold the object using 2 possible force levels (low force, LF, and high force, HF). A 10 mm x 10 mm square of 4 red light-emitting diodes (LEDs) and one yellow LED at the center was used to display the two visual cues (C1 and C2) that served as task instructions. The instructions were coded as follows: the bottom two LEDs coded for LF, the top two for HF, the leftmost two for SG, and the rightmost two for PG. By definition, PG was obtained by placing the tips of the index and the thumb in a groove on the upper and lower sides of the object, respectively, and SG by placing the tip of the thumb and the lateral surface of the index on the right and left sides, respectively. An electromagnet placed inside the apparatus was used to change the object’s effective weight (100g or 200g) to require a LF (magnet off) or HF (magnet on), respectively.

Behavioral stability was ensured by requiring animals to hold on a pressure-sensitive platform throughout the preparatory period. The animal needed to exert a relatively high force (~1.3 N) to keep the platform deflected down. Trials in which the pressure on the platform was released before the Go signal were automatically aborted.

### Trial structure

The structure of a trial was as follows: the animal started from a home position with their working hand pressing down on a pressure-sensitive switch. After a fixed 400-ms delay, the central LED was illuminated to indicate the start of a new trial. After another 400 ms, the first instruction (C1) was presented for 300 ms, followed by a 1-s preparatory period with only the central LED on. At the end of the preparatory period, the second instruction (C2) was presented and also served as the imperative GO signal. At that point, the animal needed to (1) release the switch, (2) reach for the object with the appropriate grip and (3) pull and hold it with the appropriate force for 500 ms before receiving a reward (mixture of apple sauce and water). The trial was aborted and unrewarded if the monkey released the home-switch before the GO signal or used the wrong grip type to grasp the object. To initiate a new trial, the monkey had to return their working hand to the home position and press the switch.

### Experimental conditions

The task included 4 trial types (2 grips x 2 forces), namely SG-LF, SG-HF, PG-LF and PG-HF, which were randomly interleaved across trials. To manipulate movement preparation in this task, we varied the order in which the grip and force information were provided. Specifically, in the “grip-cued” condition, C1 provided the grip instruction, while C2 provided the force instruction. Conversely, in the “grip-uncued” condition, C1 provided the force instruction, while C2 provided the grip instruction. As a result, instructed movements were identical in the grip-cued and grip-uncued trials, but differed in the order that the grip/force information necessary to plan the movement was given. Animals performed the task in short sessions of 12 to 15 minutes (blocks of uninterrupted trials) so as to get a minimum of 10 trials for each trial type. The two conditions were alternated between sessions.

### Neural recordings

Spiking data were recorded during multiple behavioral sessions (n=8 for monkey N, n=24 for monkey L). Each session had on average n=89 simultaneously recorded neurons, and n=142 successful trials. Spikes were sorted offline using Offline Spike Sorter, version 3, Plexon Inc., Dallas, TX, USA). Spike clusters which were separated significantly from each other and with less than 1% of inter-spike intervals (ISIs) of 2 ms and less were considered as single units (single-unit activity, SUA), whereas less well separated clusters and/or more than 1% of 2 ms ISIs were considered as multi-unit (multi-unit activity, MUA) recordings. In all analyses, we included both SUA and MUA.

### Analysis of neural activity

#### Neural tuning analysis

To assess the tuning properties of individual neurons, we compute a grip tuning depth of each neuron (Figure 2B). By definition, the tuning depth was given by the difference 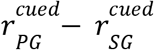, where 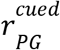 and 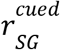 designate the firing rate in the PG-cued and SG-cued condition, respectively. To compute firing rates (see also Figure 2A), we smoothed trial-averaged spike counts in 20-ms bins using a Gaussian kernel (SD = 40 ms). We obtained confidence intervals on firing rates via standard bootstrapping by randomly sampling trials with replacement (N=100 repeats).

To quantify the relationship between firing rates in the grip-cued and grip-uncued conditions, we computed the index 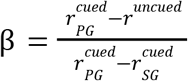, where *r^uncued^* designates the firing rate in the grip-uncued condition associated with either of the two force levels (HF or LF). This ratio was computed only for grip-selective neurons, i.e., neurons whose firing rates differed significantly (as assessed by non-overlapping 95% confidence intervals) between PG-cued and SG-cued conditions.

In Figure 2D, we computed *β* using activity in a 20-ms bin centered at two different time points of the trial: 200 ms before Go, and 400 ms after Go. Before Go, we treated the two force conditions separately, and pooled them to plot the distribution of across *β* neurons. After Go, we treated the PG-cued and SG-cued conditions separately.

#### Neural distance analysis

To quantify the strength of grip and force encoding (Figure S2A), we computed the distance in the state space between the neural trajectories associated with the PG-cued and SG-cued conditions, and between HF-cued and LF-cued. We followed the method developed in (Willett et al., 2020) to obtain an unbiased estimate of the (squared) distance by computing the squared distance based on the jackknife (leave-one-out) method across trials to calculate a 95% CI (shaded area); see further details at https://github.com/fwillett/cvVectorStats. This cross-validated approach is more computationally involved but guarantees that the distance estimate reaches zero despite single trial variability within conditions when neural trajectories overlap. We further normalized the distance separating the trajectories by the average squared distance from every time point of the PG-cued trajectory (or HF-cued) to its centroid as a way of normalizing by the trajectory “radius”.

#### Dimensionality reduction and projection onto the grip dimension

To evaluate the relationship between the population state in the grip-uncued condition relative to the two grip-cued conditions, we defined the “grip dimension” which separated the PG-cued and SG-cued trajectories. The grip dimension was defined at each time point of the trial as the unit vector connecting the SG-cued (trial-averaged) state to the PG-cued (trial-averaged) state. In Figure 3B and S2B, we projected the trial-averaged neural state binned at 20-ms and smoothed with a Gaussian kernel (SD=40ms) associated with each condition (PG-cued, SG-cued and grip-uncued) onto the grip dimension to assess the proximity of the grip-uncued state to the other two states. The grip-uncued trajectory was always defined in a force-specific manner, i.e., not mixing HF and LF trials. To obtain confidence intervals, we used standard bootstrapping (resampling trials with replacement), with 100 repeats.

In Figure 4 and S3A, we projected single trials rather than trial-averaged activity onto the grip dimension. However, the grip dimension was still defined as before using trial-averaged states associated with the PG-cued and SG-cued conditions. Single-trial activity in the grip-uncued condition was computed using a 200-ms sliding window (shifted by 1-ms steps to obtain smooth trajectories).

#### Calculation of the neural and behavioral contrast

In Figure 3D and S4B, we used two separate contrast measures (λ_*neural*_ and λ_*behavioral*_) to assess the relationship between preparatory states and reaction times on a session-by-session level. By definition, the neural contrast was given by: 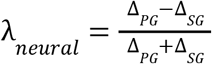, where Δ_*PG*_ and Δ_*SG*_ respectively represent the distance of the grip-uncued projection to the PG-cued and SG-cued state. Mathematically, these distances were computed by taking the absolute difference between the (scalar) projection of the cued and uncued state along the grip dimension. The neural contrast was computed for each recording session separately. Similarly, the behavioral contrast was computed for each session and defined as: 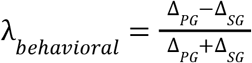, where Δ_*PG*_ and Δ_*SG*_ respectively represent the (absolute) mean reaction time difference between the grip-uncued and grip-cued condition for PG and SG. Both contrast measures, by construction, were between −1 and 1.

To compute 95% confidence intervals for the correlation values between λ_*neural*_ and λ_*behavioral*_ (shaded area in Figure 3E), we proceeded as follows: we first applied the Fisher’s transformation 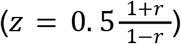 to the correlation value *r*. We then calculated the CI as: [*z* − *z_critical_ SE*; *z* + *z_critical_ SE*], where *z_critical_* = 1.96 (corresponding to 95% confidence) and *SE* is the standard error 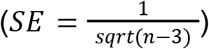. We finally transposed *z* back to correlation values using the formula: *r* = *tanh*(*z*).

#### Control analysis for the correlation between contrast measures

To assess the specificity of the relationship between the neural projections and reaction times with respect to the grip dimension, we performed a control analysis where we projected the neural states onto a random dimension. To allow for a fair comparison, we chose a dimension which was orthogonal to the grip dimension but captured as much variance as possible in the neural data. To do so, we proceeded as follows: we applied PCA to the time-varying dynamics of the PG-cued condition after projecting out the grip dimension. This allowed us to work in the null space of the grip dimension, as suggested by the reviewer. We then picked as the null dimension the first principal component to capture maximum variance.

#### Quantification of neural “vacillations”

In Figure 5B, we compute the likelihood that individual grip-uncued trials “belonged” to the PG-cued or SG-cued condition. For this analysis, we binned single-trial spiking activity in a 100-ms sliding window (shifted by a 1-ms step to obtain smooth results). For every time step, we defined the grip dimension as usual using trial-averaged activity. We then projected individual trials of the PG-cued, SG-cued and grip-uncued conditions onto the grip dimension. We then fitted a Gaussian distribution (“normfit” in Matlab) to the PG-cued and SG-cued distributions. Finally, for every grip-uncued trial, we computed the likelihoods that the associated projection belonged to the PG-cued or the SG-cued fitted distribution. To interpret the result, we computed the ratio of the PG and SG likelihoods, and took the log10 of this ratio to obtain the log likelihood ratio.

## Acknowledgements

NM was funded by a postdoctoral fellowship from the European Molecular Biology Organization (EMBO ALTF 329-2021). TB and AR were supported by the RIKEN-CNRS Collaborative Research Agreement, the Helmholtz portfolio theme “Supercomputing and modeling for the human brain” (SMHB), and the ANR-GRASP (France). The authors wish to thank Manuel Zaepffel and Margaux Duret for participating in the data collection. The authors also thank the three anonymous reviewers for their constructive feedback.

## Author contributions

TB and AR conceived the project. AR processed the behavioral and electrophysiological data. NM performed all the analyses and wrote the manuscript. All authors contributed to the interpretation of the data. TB supervised the project.

## Data and code availability

The data and code used in the study will be made available upon publication.

## Supplementary material

**Table S1.**
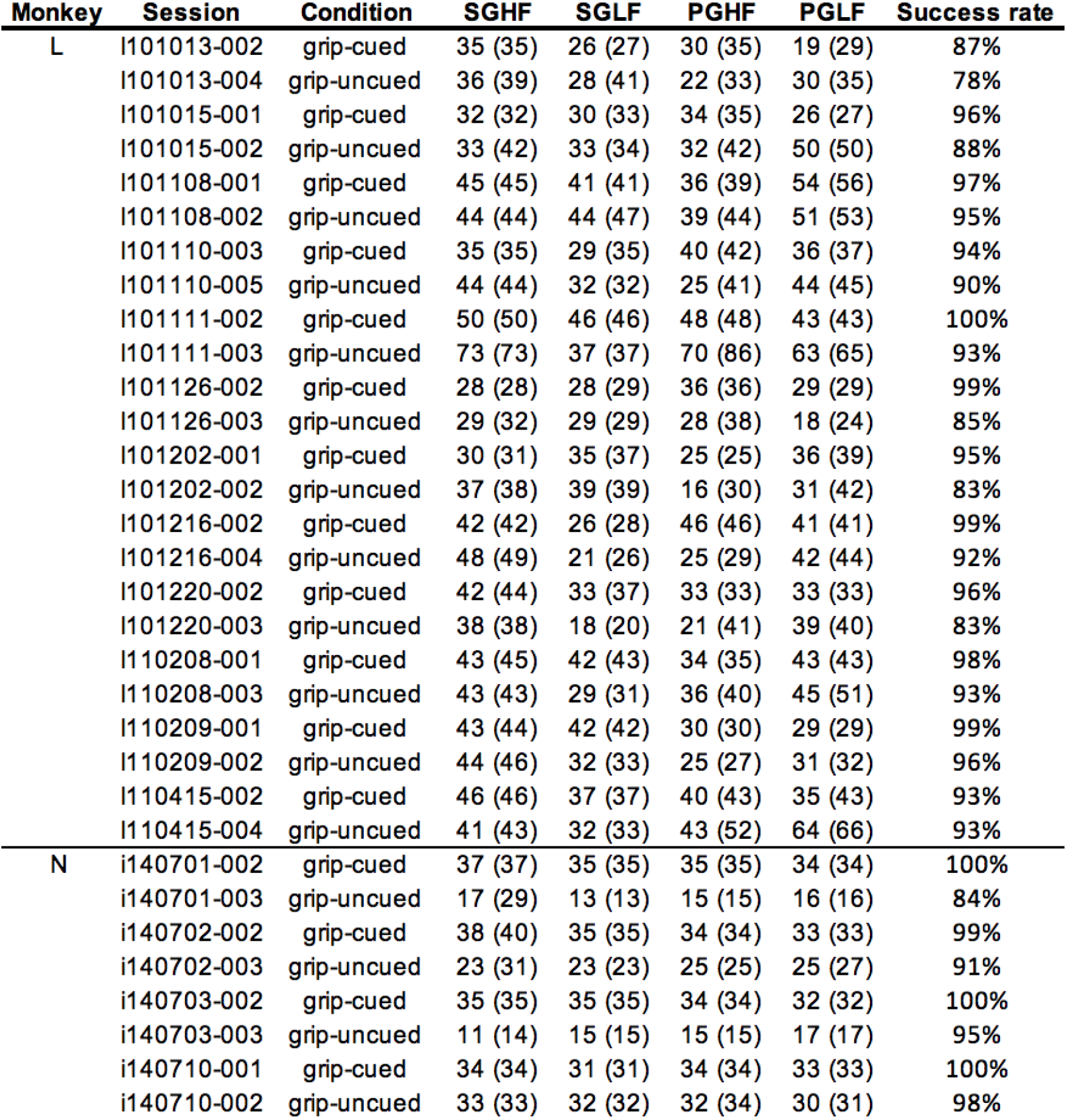
Trial statistics and success rates for individual sessions and conditions for the two animals (n=24 sessions for monkey L, 8 for monkey N). The number shown in each trial type column corresponds to the number of successful trials, followed by the total number of trials in parentheses. Success rates are averaged across all trial types of each condition.

**Figure S1.**
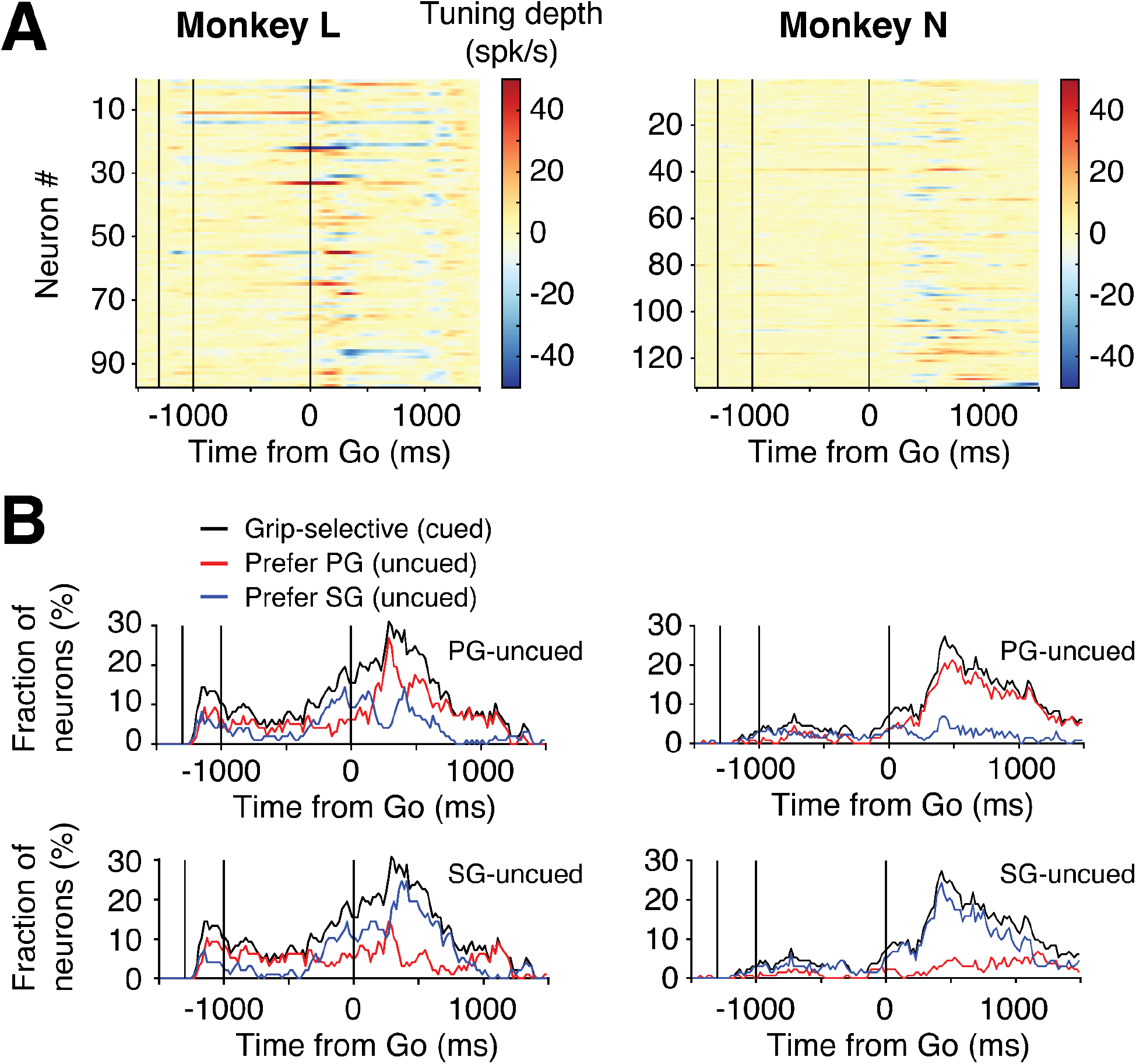
Related to Figure 2. **(A)** Grip tuning depth across neurons, as measured by the instantaneous firing rate difference between the PG-cued and SG-cued activity. Each row represents one neuron, and the color indicates the magnitude of the tuning depth (blue for negative values, i.e., neuron has a larger firing rate in SG; red for positive values). **(B)** Fraction of grip-selective neurons in grip-cued condition (black line), and neurons that prefer PG (red line) or SG (blue line) in the grip-uncued condition. We considered a neuron to be grip-selective if its instantaneous firing rate in the PG-cued and SG-cued conditions were significantly different (non-overlapping 95% confidence intervals obtained via standard bootstrapping across trials). Among the grip-selective neurons, we identified neurons which preferred PG or SG in the grip-uncued condition at every time point of the trial. By definition, a neuron was considered to prefer PG (resp., SG) if its instantaneous firing rate in the grip-uncued condition was closer to its firing rate in PG-cued (resp., SG-cued).

**Figure S2.**
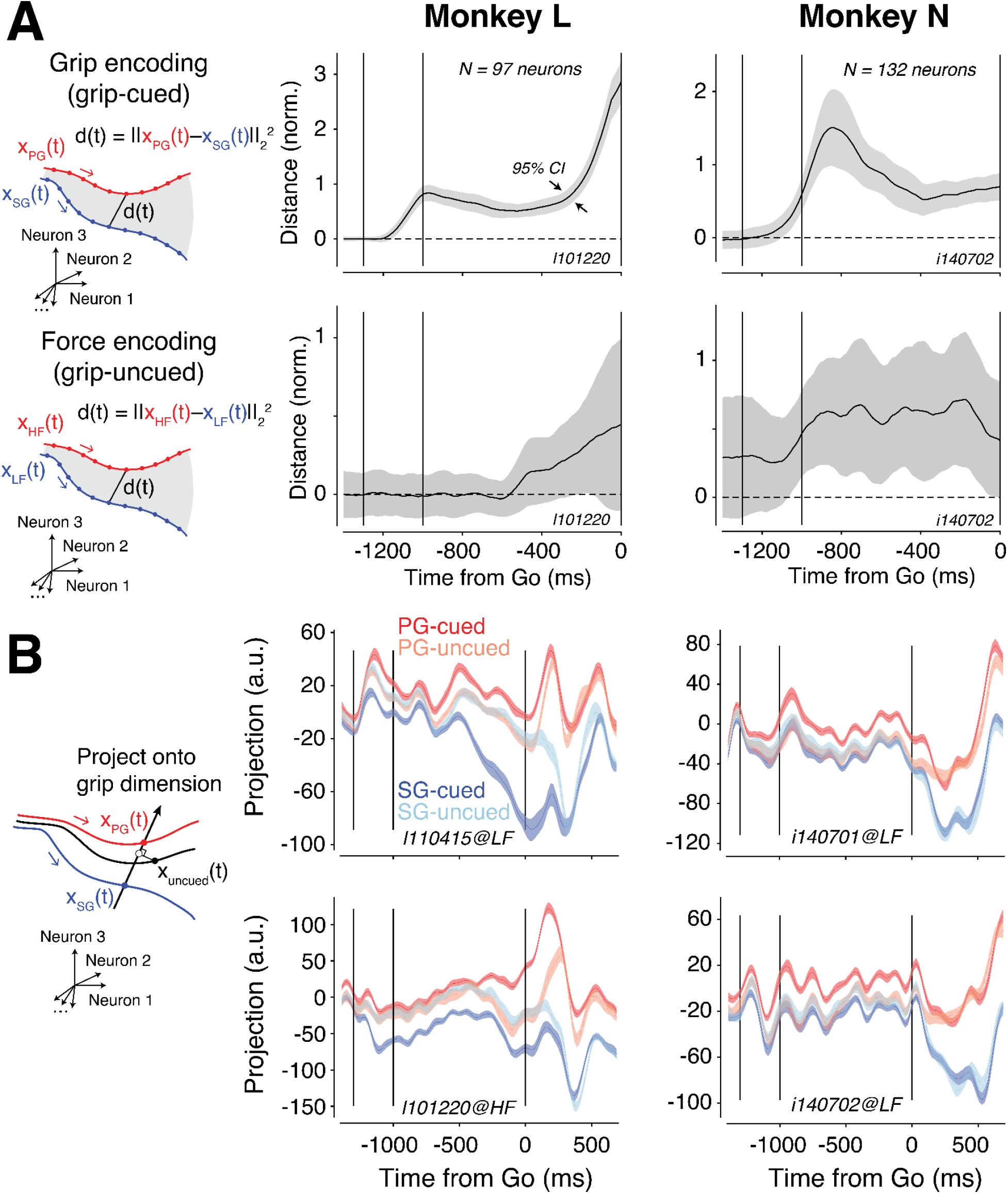
Related to Figure 3. **(A)** Comparison of grip and force encoding as measured by the distance between neural trajectories during the preparatory period. Following (Willett et al.,2020), we obtained an unbiased estimate of the distance by computing the squared distance based on the jackknife (leave-one-out) method across trials to calculate a 95% CI (shaded area); see further details in Methods. This approach guarantees that the distance estimate reaches zero despite single trial variability within conditions when neural trajectories overlap. We further normalized the result by the average squared distance from every time point of the PG-cued trajectory to its centroid (as a measure of the trajectory “radius”) to allow comparison across animals. Note the slight difference in the distance profile (grip-cued; top) between the two monkeys. We attribute this difference to the fact that monkey N did not anticipate the Go signal as much as monkey L, as evidenced by longer reaction times (Figure 1C). **(B)** Projection of the neural trajectories associated with the PG-cued (dark red), SG-cued (dark blue), PG-uncued (light red), and SG-uncued (light blue) condition onto the grip dimension for different sessions and animals. For the PG-uncued and SG-uncued conditions, we fixed the level of force (HF for bottom left panel, LF for other panels). The slight separation of PG-cued and SG-cued projections before the Cue onset is an artifact of defining the grip dimension in a non-cross-validated way (which is not necessary for this particular analysis, since we are interested in projecting grip-uncued trials, which by definition are cross-validated). Note however that a cross-validated neural distance analysis confirmed that the PG-cued and SG-cued trajectories overlapped prior to Cue onset (see Figure S2A).

**Figure S3.**
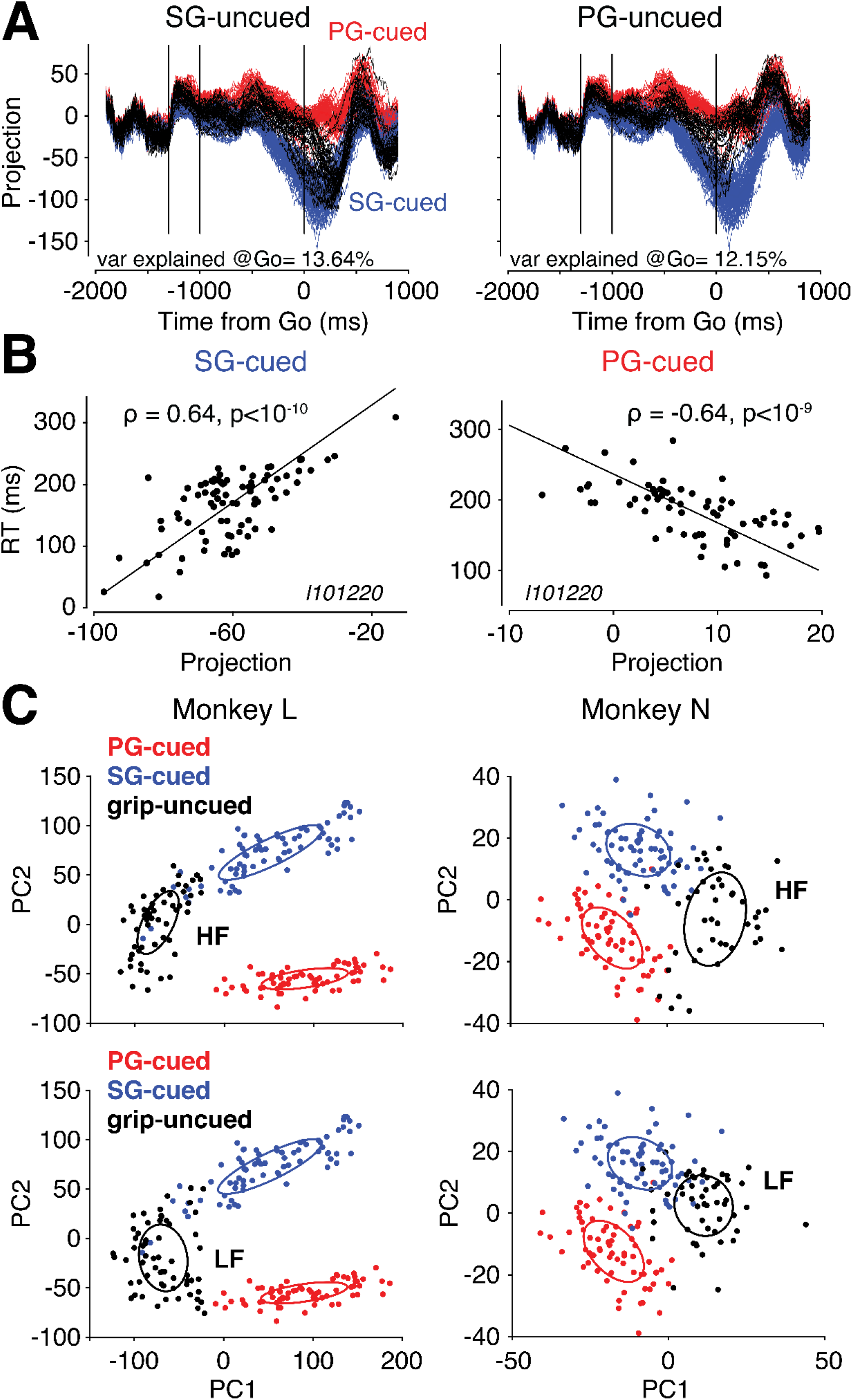
Related to Figure 4. **(A)** Single-trial projections of SG-uncued (left) and PG-uncued onto the grip dimension. Each black line represents a single trial of the grip-uncued condition (fixed at HF); red and blue lines represent single trials of the PG-cued and SG-cued condition. Single-trial activity was computed using a 200-ms sliding window to bin the spikes. The amount of variance across trials at the time of Go (200-ms window preceding Go) along the grip dimension is indicated at the bottom of each panel. By comparison, the amount of variance along a random direction was around 1%. **(B)** Correlation between reaction time and single-trial projections onto the grip dimension right before Go for the SG-cued (left) and PG-cued (right) condition. **(C)** Two-dimensional projections of neural states at the time of Go for all conditions (red for PG-cued; blue for SG-cued; black for grip-uncued, HF at top and LF at bottom) for both monkeys. Principal component analysis was performed on the trial-averaged preparatory states across all four conditions. Each individual dot represents a single trial. Ellipses are centered around the condition-average and scaled by the standard deviation across trials.

**Figure S4.**
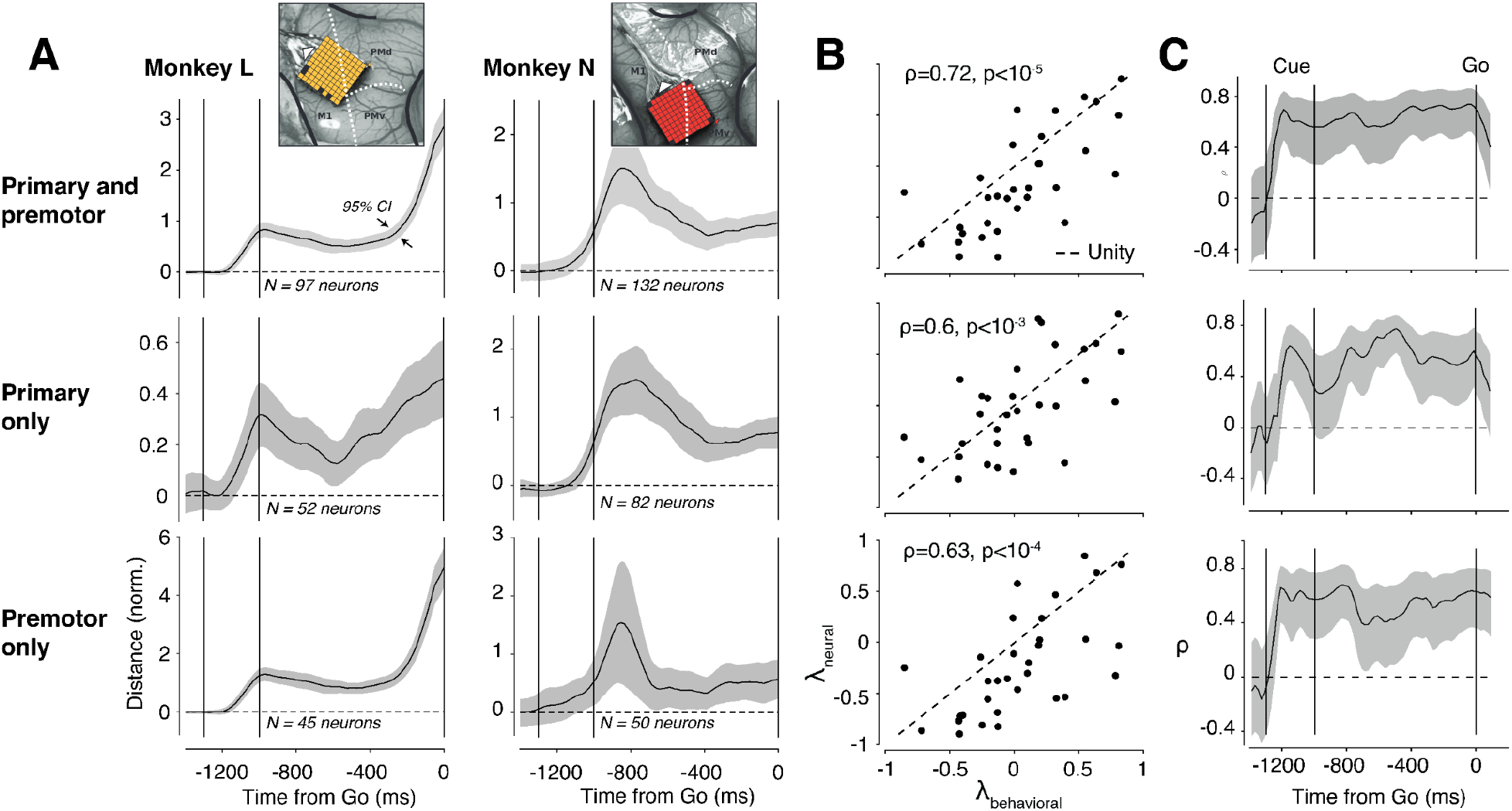
Related to Figure 3. **(A)** Distance between neural trajectories associated with the PG-cued and SG-cued conditions (similar to Figure S2A). At the top is shown the location of the Utah array for each monkey. **(B)** Correlation between neural projections along the grip dimension and reaction times (similar to Figure 3D). **(C)** Evolution of the correlation between neural projections and reaction times as a function of time in the trial (similar to Figure 3E). In all three panels, we either used all neurons (top; M1+PM), only neurons from the posterior (middle; M1) or anterior (bottom; PM) half of the Utah array.

**Figure S5.**
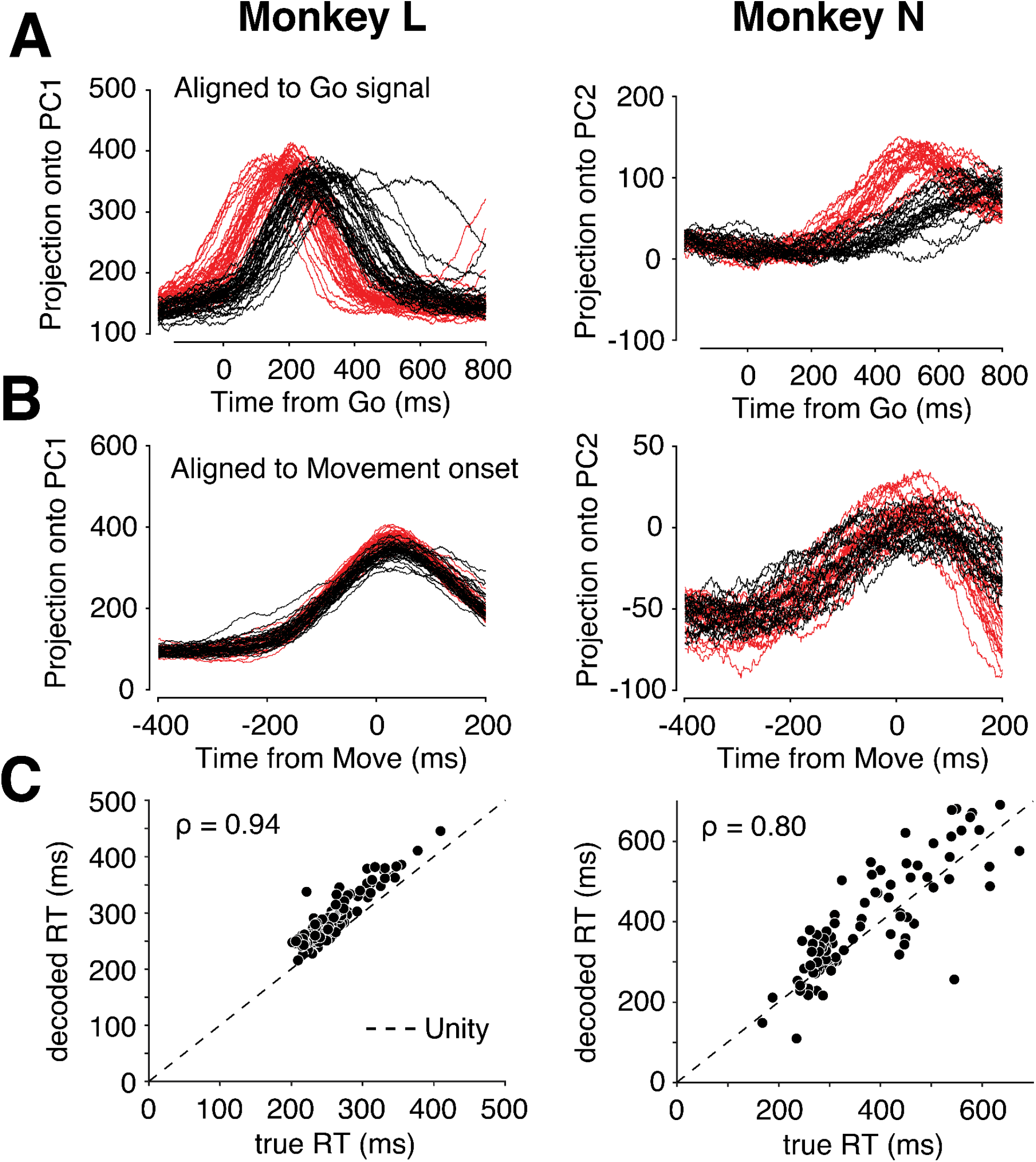
Related to Figure 4. **(A)** One-dimensional projection of single-trial neural dynamics around movement initiation. Data is aligned to the Go signal. We overlaid dynamics of the grip-cued (red traces) and grip-uncued (black traces) conditions. To avoid clutter, only trials corresponding to PG-LF/LF-PG movements are shown for both monkeys (other trial types looked qualitatively similar). To enforce cross-validation, the 1-D projections were obtained after applying principal component analysis only to the grip-cued trial-averaged dynamics (between Go-300ms and Go+900ms), and then projecting grip-cued and grip-uncued single-trial dynamics onto the top principal component. For monkey N, single-trial projections along PC2 were less noisy so we used that component to describe the dynamics. For this analysis, spike trains were binned in a 200-ms sliding window. **(B)** Same as (A) but the data is aligned to Move (i.e., movement onset). The window used for PCA was defined between Move-500ms and Move+300ms. **(C)** Ground truth reaction times versus decoded reaction times based on a threshold-crossing hypothesis. For each trial type (e.g., PG-HF/HF-PG), we defined a threshold value in a cross-validated way as follows: for each trial, we computed the time that the dynamics approached closest to a set threshold value, and considered this time as the “decoded” reaction time on that trial. We then optimized the value of the threshold to minimize the RMSE between the true and the decoded reaction times across grip-cued trials only. Using the same threshold value, we then computed the time that the dynamics in the grip-uncued condition approached closest to the threshold, and considered this time as the (cross-validated) decoded reaction time. In the plot, only the grip-uncued trials are shown, and combined across the 4 trial types. The correlation value at the top indicates the strength of the correlation between the decoded and the true reaction times across trials.

## References

Afshar, A., Santhanam, G., Yu, B. M., Ryu, S. I., Sahani, M., & Shenoy, K. V. (2011). Single-Trial Neural Correlates of Arm Movement Preparation. Neuron, 71(3), 555–564.

Alhussein, L., & Smith, M. A. (2021). Motor planning under uncertainty. eLife, 10. https://doi.org/10.7554/eLife.67019

Ames, K. C., Ryu, S. I., & Shenoy, K. V. (2014). Neural Dynamics of Reaching following Incorrect or Absent Motor Preparation. Neuron, 81(2), 438–451.

Ames, K. C., Ryu, S. I., & Shenoy, K. V. (2019). Simultaneous motor preparation and execution in a last-moment reach correction task. Nature Communications, 10(1), 2718.

Arai, K., McPeek, R. M., & Keller, E. L. (2004). Properties of saccadic responses in monkey when multiple competing visual stimuli are present. Journal of Neurophysiology, 91(2), 890–900.

Athalye, V. R., Carmena, J. M., & Costa, R. M. (2020). Neural reinforcement: re-entering and refining neural dynamics leading to desirable outcomes. Current Opinion in Neurobiology, 60, 145–154.

Bastian, A., Schöner, G., & Riehle, A. (2003). Preshaping and continuous evolution of motor cortical representations during movement preparation. The European Journal of Neuroscience, 18(7), 2047–2058.

Batista, A. P., Santhanam, G., Yu, B. M., Ryu, S. I., Afshar, A., & Shenoy, K. V. (2007). Reference frames for reach planning in macaque dorsal premotor cortex. Journal of Neurophysiology, 98(2), 966–983.

Baumann, M. A., Fluet, M.-C., & Scherberger, H. (2009). Context-specific grasp movement representation in the macaque anterior intraparietal area. The Journal of Neuroscience: The Official Journal of the Society for Neuroscience, 29(20), 6436–6448.

Beiran, M., Meirhaeghe, N., Sohn, H., Jazayeri, M., & Ostojic, S. (2021). Parametric control of flexible timing through low-dimensional neural manifolds. In bioRxiv (p.2021.11.08.467806). https://doi.org/10.1101/2021.11.08.467806

Brochier, T., Zehl, L., Hao, Y., Duret, M., Sprenger, J., Denker, M., Grün, S., & Riehle, A. (2018). Massively parallel recordings in macaque motor cortex during an instructed delayed reach-to-grasp task. Scientific Data, 5, 180055.

Chapman, C. S., Gallivan, J. P., Wood, D. K., Milne, J. L., Culham, J. C., & Goodale, M. A. (2010). Reaching for the unknown: Multiple target encoding and real-time decision-making in a rapid reach task. Cognition, 116(2), 168–176.

Chou, I. H., Sommer, M. A., & Schiller, P. H. (1999). Express averaging saccades in monkeys. Vision Research, 39(25), 4200–4216.

Churchland, M. M., Cunningham, J. P., Kaufman, M. T., Foster, J. D., Nuyujukian, P., Ryu, S. I., & Shenoy, K. V. (2012). Neural population dynamics during reaching. Nature, 487, 51.

Churchland, M. M., Cunningham, J. P., Kaufman, M. T., Ryu, S. I., & Shenoy, K. V. (2010). Cortical preparatory activity: representation of movement or first cog in a dynamical machine? Neuron, 68(3), 387–400.

Churchland, M. M., Santhanam, G., & Shenoy, K. V. (2006). Preparatory activity in premotor and motor cortex reflects the speed of the upcoming reach. Journal of Neurophysiology, 96(6), 3130–3146.

Churchland, M. M., & Shenoy, K. V. (2007a). Delay of movement caused by disruption of cortical preparatory activity. Journal of Neurophysiology, 97(1), 348–359.

Churchland, M. M., & Shenoy, K. V. (2007b). Temporal complexity and heterogeneity of single-neuron activity in premotor and motor cortex. Journal of Neurophysiology, 97(6), 4235–4257.

Churchland, M. M., Yu, B. M., Ryu, S. I., Santhanam, G., & Shenoy, K. V. (2006). Neural variability in premotor cortex provides a signature of motor preparation. The Journal of Neuroscience: The Official Journal of the Society for Neuroscience, 26(14), 3697–3712.

Cisek, P. (2006). Integrated neural processes for defining potential actions and deciding between them: a computational model. The Journal of Neuroscience: The Official Journal of the Society for Neuroscience, 26(38), 9761–9770.

Cisek, P. (2007). Cortical mechanisms of action selection: the affordance competition hypothesis. Philosophical Transactions of the Royal Society of London. Series B,Biological Sciences, 362(1485), 1585–1599.

Cisek, P. (2012). Making decisions through a distributed consensus. Current Opinion in Neurobiology, 22(6), 927–936.

Cisek, P., & Kalaska, J. F. (2005). Neural correlates of reaching decisions in dorsal premotor cortex: specification of multiple direction choices and final selection of action. Neuron, 45(5), 801–814.

Coallier, É., Michelet, T., & Kalaska, J. F. (2015). Dorsal premotor cortex: neural correlates of reach target decisions based on a color-location matching rule and conflicting sensory evidence. Journal of Neurophysiology, 113(10), 3543–3573.

Costello, M. G., Zhu, D., Salinas, E., & Stanford, T. R. (2013). Perceptual modulation of motor--but not visual--responses in the frontal eye field during an urgent-decision task. The Journal of Neuroscience: The Official Journal of the Society for Neuroscience, 33(41), 16394–16408.

Crammond, D. J., & Kalaska, J. F. (2000). Prior information in motor and premotor cortex: activity during the delay period and effect on pre-movement activity. Journal of Neurophysiology, 84(2), 986–1005.

Day, B. L., Rothwell, J. C., Thompson, P. D., Maertens de Noordhout, A., Nakashima, K., Shannon, K., & Marsden, C. D. (1989). Delay in the execution of voluntary movement by electrical or magnetic brain stimulation in intact man. Evidence for the storage of motor programs in the brain. Brain: A Journal of Neurology, 112 (Pt3), 649–663.

Dekleva, B. M., Kording, K. P., & Miller, L. E. (2018). Single reach plans in dorsal premotor cortex during a two-target task. Nature Communications, 9(1), 3556.

Dekleva, B. M., Ramkumar, P., Wanda, P. A., Kording, K. P., & Miller, L. E. (2016). Uncertainty leads to persistent effects on reach representations in dorsal premotor cortex. eLife, 5. https://doi.org/10.7554/eLife.14316

Denker, M., Zehl, L., Kilavik, B. E., Diesmann, M., Brochier, T., Riehle, A., & Grün, S. (2018). LFP beta amplitude is linked to mesoscopic spatio-temporal phase patterns. Scientific Reports, 8(1), 5200.

Dubreuil, A., Valente, A., Beiran, M., Mastrogiuseppe, F., & Ostojic, S. (2022). The role of population structure in computations through neural dynamics. Nature Neuroscience, 1–12.

Elsayed, G. F., Lara, A. H., Kaufman, M. T., Churchland, M. M., & Cunningham, J. P. (2016). Reorganization between preparatory and movement population responses in motor cortex. Nature Communications, 7, 13239.

Erlhagen, W., & Schöner, G. (2002). Dynamic field theory of movement preparation. Psychological Review, 109(3), 545–572.

Even-Chen, N., Sheffer, B., Vyas, S., Ryu, S. I., & Shenoy, K. V. (2019). Structure and variability of delay activity in premotor cortex. PLoS Computational Biology, 15(2), e1006808.

Fetz, E. E. (1992). Are movement parameters recognizably coded in the activity of single neurons? The Behavioral and Brain Sciences, 15(4), 679–690.

Fluet, M.-C., Baumann, M. A., & Scherberger, H. (2010). Context-specific grasp movement representation in macaque ventral premotor cortex. The Journal of Neuroscience: The Official Journal of the Society for Neuroscience, 30(45), 15175–15184.

Gallivan, J. P., Barton, K. S., Chapman, C. S., Wolpert, D. M., & Flanagan, J. R. (2015). Action plan co-optimization reveals the parallel encoding of competing reach movements. Nature Communications, 6, 7428.

Gallivan, J. P., & Chapman, C. S. (2014). Three-dimensional reach trajectories as a probe of real-time decision-making between multiple competing targets. Frontiers in Neuroscience, 8, 215.

Ghez, C., Favilla, M., Ghilardi, M. F., Gordon, J., Bermejo, R., & Pullman, S. (1997). Discrete and continuous planning of hand movements and isometric force trajectories. Experimental Brain Research. Experimentelle Hirnforschung.Experimentation Cerebrale, 115(2), 217–233.

Godschalk, M., Lemon, R. N., Kuypers, H. G., & van der Steen, J. (1985). The involvement of monkey premotor cortex neurones in preparation of visually cued arm movements. Behavioural Brain Research, 18(2), 143–157.

Golub, M. D., Sadtler, P. T., Oby, E. R., Quick, K. M., Ryu, S. I., Tyler-Kabara, E. C., Batista, A. P., Chase, S. M., & Yu, B. M. (2018). Learning by neural reassociation. Nature Neuroscience, 21(4), 607–616.

Haith, A. M., Huberdeau, D. M., & Krakauer, J. W. (2015). Hedging your bets: intermediate movements as optimal behavior in the context of an incomplete decision. PLoS Computational Biology, 11(3), e1004171.

Hatsopoulos, N. G., Xu, Q., & Amit, Y. (2007). Encoding of movement fragments in the motor cortex. The Journal of Neuroscience: The Official Journal of the Society for Neuroscience, 27(19), 5105–5114.

Hennequin, G., Vogels, T. P., & Gerstner, W. (2014). Optimal Control of Transient Dynamics in Balanced Networks Supports Generation of Complex Movements. Neuron, 82(6), 1394–1406.

Hudson, T. E., Maloney, L. T., & Landy, M. S. (2007). Movement planning with probabilistic target information. Journal of Neurophysiology, 98(5), 3034–3046.

Ifft, P. J., Lebedev, M. A., & Nicolelis, M. A. L. (2012). Reprogramming movements: extraction of motor intentions from cortical ensemble activity when movement goals change. Frontiers in Neuroengineering, 5, 16.

Kalaska, J. F., Scott, S. H., Cisek, P., & Sergio, L. E. (1997). Cortical control of reaching movements. Current Opinion in Neurobiology, 7(6), 849–859.

Kao, T.-C., Sadabadi, M. S., & Hennequin, G. (2021). Optimal anticipatory control as a theory of motor preparation: A thalamo-cortical circuit model. Neuron, 109(9), 1567–1581.e12.

Kaufman, M. T., Churchland, M. M., Ryu, S. I., & Shenoy, K. V. (2014). Cortical activity in the null space: permitting preparation without movement. Nature Neuroscience, 17, 440.

Kaufman, M. T., Churchland, M. M., Ryu, S. I., & Shenoy, K. V. (2015). Vacillation,indecision and hesitation in moment-by-moment decoding of monkey motor cortex. eLife, 4, e04677.

Klaes, C., Westendorff, S., Chakrabarti, S., & Gail, A. (2011). Choosing goals, not rules: deciding among rule-based action plans. Neuron, 70(3), 536–548.

Kurata, K. (1993). Premotor cortex of monkeys: set- and movement-related activity reflecting amplitude and direction of wrist movements. Journal of Neurophysiology, 69(1), 187–200.

Lara, A. H., Elsayed, G. F., Zimnik, A. J., Cunningham, J. P., & Churchland, M. M. (2018). Conservation of preparatory neural events in monkey motor cortex regardless of how movement is initiated. eLife, 7. https://doi.org/10.7554/eLife.31826

Logiaco, L., Abbott, L. F., & Escola, S. (2021). Thalamic control of cortical dynamics in a model of flexible motor sequencing. Cell Reports, 35(9), 109090.

Messier, J., & Kalaska, J. F. (2000). Covariation of primate dorsal premotor cell activity with direction and amplitude during a memorized-delay reaching task. Journal of Neurophysiology, 84(1), 152–165.

Michaels, J. A., Dann, B., Intveld, R. W., & Scherberger, H. (2015). Predicting Reaction Time from the Neural State Space of the Premotor and Parietal Grasping Network. The Journal of Neuroscience: The Official Journal of the Society for Neuroscience, 35(32), 11415–11432.

Milekovic, T., Truccolo, W., Grün, S., Riehle, A., & Brochier, T. (2015). Local field potentials in primate motor cortex encode grasp kinetic parameters. NeuroImage, 114, 338–355.

Pandarinath, C., O’Shea, D. J., Collins, J., Jozefowicz, R., Stavisky, S. D., Kao, J. C., Trautmann, E. M., Kaufman, M. T., Ryu, S. I., Hochberg, L. R., Henderson, J. M., Shenoy, K. V., Abbott, L. F., & Sussillo, D. (2018). Inferring single-trial neural population dynamics using sequential auto-encoders. Nature Methods, 15(10),805–815.

Peixoto, D., Verhein, J. R., Kiani, R., Kao, J. C., Nuyujukian, P., Chandrasekaran, C., Brown, J., Fong, S., Ryu, S. I., Shenoy, K. V., & Newsome, W. T. (2021). Decoding and perturbing decision states in real time. Nature, 591(7851), 604–609.

Remington, E. D., Narain, D., Hosseini, E. A., & Jazayeri, M. (2018). Flexible Sensorimotor Computations through Rapid Reconfiguration of Cortical Dynamics. Neuron, 98(5), 1005–1019.e5.

Requin, J., Brener, J., & Ring, C. (1991). Preparation for action. Handbook of Cognitive Psychophysiology: Central and Autonomic Nervous System Approaches., 745, 357–448.

Rickert, J., Riehle, A., Aertsen, A., Rotter, S., & Nawrot, M. P. (2009). Dynamic encoding of movement direction in motor cortical neurons. The Journal of Neuroscience: The Official Journal of the Society for Neuroscience, 29(44), 13870–13882.

Riehle, A. (2005). Preparation for action: one of the key functions of the motor cortex. In Riehle, A. & Vaadia, E. (Eds), Motor Cortex in Voluntary Movements: A Distributed System for Distributed Functions, CRC-Press, Boca Raton, FL. http://citeseerx.ist.psu.edu/viewdoc/summary?doi=10.1.1.486.7682

Riehle, A., Brochier, T., Nawrot, M., & Grün, S. (2018). Behavioral Context Determines Network State and Variability Dynamics in Monkey Motor Cortex. Frontiers in Neural Circuits, 12, 52.

Riehle, A., MacKay, W. A., & Requin, J. (1994). Are extent and force independent movement parameters? Preparation-and movement-related neuronal activity in the monkey cortex. Experimental Brain Research. Experimentelle Hirnforschung.Experimentation Cerebrale, 99(1), 56–74.

Riehle, A., & Requin, J. (1989). Monkey primary motor and premotor cortex: single-cell activity related to prior information about direction and extent of an intended movement. Journal of Neurophysiology, 61(3), 534–549.

Riehle, A., & Requin, J. (1993). The predictive value for performance speed of preparatory changes in neuronal activity of the monkey motor and premotor cortex. Behavioural Brain Research, 53(1-2), 35–49.

Riehle, A., Wirtssohn, S., Grün, S., & Brochier, T. (2013). Mapping the spatio-temporal structure of motor cortical LFP and spiking activities during reach-to-grasp movements. Frontiers in Neural Circuits, 7, 48.

Rosenbaum, D. A. (1980). Human movement initiation: Specification of arm, direction, and extent. Journal of Experimental Psychology. General, 109(4), 444–474.

Sadtler, P. T., Quick, K. M., Golub, M. D., Chase, S. M., Ryu, S. I., Tyler-Kabara, E. C., Yu, B. M., & Batista, A. P. (2014). Neural constraints on learning. Nature, 512(7515), 423–426.

Sauerbrei, B. A., Guo, J.-Z., Cohen, J. D., Mischiati, M., Guo, W., Kabra, M., Verma, N., Mensh, B., Branson, K., & Hantman, A. W. (2020). Cortical pattern generation during dexterous movement is input-driven. Nature, 577(7790), 386–391.

Saxena, S., & Cunningham, J. P. (2019). Towards the neural population doctrine. Current Opinion in Neurobiology, 55, 103–111.

Scott, S. H. (2008). Inconvenient truths about neural processing in primary motor cortex. The Journal of Physiology, 586(5), 1217–1224.

Shen, L., & Alexander, G. E. (1997). Preferential representation of instructed target location versus limb trajectory in dorsal premotor area. Journal of Neurophysiology, 77(3), 1195–1212.

Shenoy, K. V., Kaufman, M. T., Sahani, M., & Churchland, M. M. (2011). A dynamical systems view of motor preparation: implications for neural prosthetic system design. Progress in Brain Research, 192, 33–58.

Shenoy, K. V., Sahani, M., & Churchland, M. M. (2013). Cortical control of arm movements: a dynamical systems perspective. Annual Review of Neuroscience, 36, 337–359.

Sohn, H., Meirhaeghe, N., Rajalingham, R., & Jazayeri, M. (2020). A network perspective on sensorimotor learning. Trends in Neurosciences. https://doi.org/10.1016/j.tins.2020.11.007

Sohn, H., Narain, D., Meirhaeghe, N., & Jazayeri, M. (2019). Bayesian Computation through Cortical Latent Dynamics. Neuron. https://doi.org/10.1016/j.neuron.2019.06.012

Stewart, B. M., Baugh, L. A., Gallivan, J. P., & Flanagan, J. R. (2013). Simultaneous encoding of the direction and orientation of potential targets during reach planning: evidence of multiple competing reach plans. Journal of Neurophysiology, 110(4), 807–816.

Stewart, B. M., Gallivan, J. P., Baugh, L. A., & Flanagan, J. R. (2014). Motor, not visual,encoding of potential reach targets. Current Biology: CB, 24(19), R953–R954.

Sun, X., O’Shea, D. J., Golub, M. D., Trautmann, E. M., Vyas, S., Ryu, S. I., & Shenoy, K. V. (2020). Skill-specific changes in cortical preparatory activity during motor learning. In bioRxiv (p. 2020.01.30.919894). https://doi.org/10.1101/2020.01.30.919894

Sussillo, D., Churchland, M. M., Kaufman, M. T., & Shenoy, K. V. (2015). A neural network that finds a naturalistic solution for the production of muscle activity. Nature Neuroscience, 18(7), 1025–1033.

Tanji, J., & Evarts, E. V. (1976). Anticipatory activity of motor cortex neurons in relation to direction of an intended movement. Journal of Neurophysiology, 39(5), 1062–1068.

Thura, D., & Cisek, P. (2014). Deliberation and commitment in the premotor and primary motor cortex during dynamic decision making. Neuron, 81(6), 1401–1416.

Torre, E., Quaglio, P., Denker, M., Brochier, T., Riehle, A., & Grün, S. (2016). Synchronous Spike Patterns in Macaque Motor Cortex during an Instructed-Delay Reach-to-Grasp Task. The Journal of Neuroscience: The Official Journal of the Society for Neuroscience, 36(32), 8329–8340.

Vyas, S., Even-Chen, N., Stavisky, S. D., Ryu, S. I., Nuyujukian, P., & Shenoy, K. V. (2018). Neural Population Dynamics Underlying Motor Learning Transfer. Neuron, 97(5), 1177–1186.e3.

Vyas, S., Golub, M. D., Sussillo, D., & Shenoy, K. V. (2020). Computation Through Neural Population Dynamics. Annual Review of Neuroscience, 43(1), 249–275.

Vyas, S., O’Shea, D. J., Ryu, S. I., & Shenoy, K. V. (2020). Causal Role of Motor Preparation during Error-Driven Learning. Neuron, 106(2), 329–339.e4.

Wang, J., Narain, D., Hosseini, E. A., & Jazayeri, M. (2017). Flexible timing by temporal scaling of cortical responses. Nature Neuroscience. https://doi.org/10.1038/s41593-017-0028-6

Weinrich, M., Wise, S. P., & Mauritz, K. H. (1984). A neurophysiological study of the premotor cortex in the rhesus monkey. Brain: A Journal of Neurology, 107(Pt 2), 385–414.

Willett, F. R., Deo, D. R., Avansino, D. T., Rezaii, P., Hochberg, L. R., Henderson, J. M., & Shenoy, K. V. (2020). Hand Knob Area of Premotor Cortex Represents the Whole Body in a Compositional Way. Cell, 181(2), 396–409.e26.

Wise, S. P. (1985). The primate premotor cortex: past, present, and preparatory. Annual Review of Neuroscience, 8, 1–19.

Wong, A. L., & Haith, A. M. (2017). Motor planning flexibly optimizes performance under uncertainty about task goals. Nature Communications, 8, 14624.

Zaepffel, M., & Brochier, T. (2012). Planning of visually guided reach-to-grasp movements: inference from reaction time and contingent negative variation (CNV). Psychophysiology, 49(1), 17–30.

Zimnik, A. J., & Churchland, M. M. (2021). Independent generation of sequence elements by motor cortex. Nature Neuroscience. https://doi.org/10.1038/s41593-021-00798-5

